# E protein control of NKγδT cell development through both generation and function of the stereotypic Vγ1Vδ6.3 TCR

**DOI:** 10.1101/2024.01.04.574274

**Authors:** Ariana Mihai, Sang-Yun Lee, Susan Shinton, Mitchell I. Parker, Alejandra V. Contreras, Baojun Zhang, Michele Rhodes, Roland L. Dunbrack, Juan-Carlos Zúñiga-Pflücker, Maria Ciofani, Yuan Zhuang, David L. Wiest

## Abstract

T cell receptor (TCR) signals regulate important developmental transitions through induction of the E protein antagonist, Id3; however, Id3-deficiency produces paradoxical effects on γδ T cell subsets. Here, we show here that Id3-deficiency attenuates the development of Vγ3-expressing γδ T cells, while markedly enhancing the development of Vγ1Vδ6.3-expressing NKγδT cells. Id3-deficiency does so by regulating both the generation of the stereotypic Vγ1Vδ6.3 TCR expressed by NKγδT cells and its capacity to support development. Indeed, we determined that the *Trav15* segment, which encodes the Vδ6.3 TCR subunit, is directly bound by E proteins that control its expression. Moreover, once expressed, the resulting Vγ1Vδ6.3 TCR in capable of specifying the innate-like NKγδT cell fate in a cell-autonomous and developmentally-unrestricted manner that is restrained by the Id3/E axis. Together, these data indicate that the paradoxical behavior of NKγδT cells in the Id3-deficient setting is entirely determined by its stereotypic Vγ1Vδ6.3 TCR complex.

## INTRODUCTION

T cells comprise two major lineages, αβ or γδ, broadly defined by the T cell antigen receptor (TCR) complex they express. These lineages arise from a common progenitor pool^1-3^, and their divergence occurs in response to differences in TCR signaling, with weak TCR signals, typically transduced by the pre-TCR complex specifying the αβ T cell fate, and stronger or more prolonged TCR signals emanating from the γδTCR and specifying the γδ T cell fate^4-6^. The TCR signals of differing strength that instruct divergence of the αβ and γδ T lineages depend on the kinases Lck and ERK, which induce transcription factors (TF) of the Egr family, and result in the proportional induction of the helix-loop-helix factor, Id3^6–8^. Id3 is a critical regulator of αβ/γδ T lineage commitment, and it functions by inhibiting DNA binding by a family of TFs termed E box DNA binding proteins (E proteins). E proteins are Class I basic helix-loop-helix (bHLH) family of TFs that bind as homo- or heterodimers to CANNTG motifs (E-box sites)^9^ and serve as critical regulators of lymphoid development and function^9–15^. The differences in TCR signal strength that control lineage fate do so by proportional induction of Id3, which produces graded reductions in E protein activity, with weak signals specifying the αβ fate through modest reduction in E protein activity and strong signals causing the profound repression of E protein activity required for γδ T lineage commitment^8,16–18^.

Importantly, Id3-deficiency was found to attenuate development of γδ T cells in a defined γδ TCR transgenic model system, supporting the model that TCR signal strength instructs the γδ T cell fate through graded repression of E protein function^8^. However, several groups identified a subset of γδ T cells, characterized by expression of a Vγ1Vδ6 TCR, that was markedly expanded in non-Tg Id3-deficient mice^8,19–21^, raising the question of why Id3-deficiency did not attenuate their development. A distinguishing aspect of γδ T lymphocyte ontogeny is the ordered appearance of waves of γδ T cells defined by their Vγ usage ^22,23^, with each wave associated with an effector function and anatomic location^24^. The first such wave, peaks around embryonic day 14 (E14) and comprises Vγ3^+^ dendritic epidermal T cells (DETCs), which produce interferon-γ (IFNγ) and home to the epidermis ^25^. The next waves occur at E16 and E18 and comprise Vγ4^+^ and Vγ2^+^ subsets, respectively, which are associated with IL-17 production and residence in the lung, uterus, and lymphoid tissues^24,26,27^. The final wave begins during late fetal life and comprises Vγ1^+^ T lymphocytes, among which are the natural killer γδ T cell (NKγδT) subset^27–30^. NKγδT cells are characterized by a strereotypic Vγ1.1Vδ6.3 TCR, PLZF expression, and co-produciton of IL-4 and IFNγ without the need for prior activation^19,31,32^; and their development is restricted to the late fetal/perinatal developmental window^19,31,32^.

Development of PLZF-expressing NKγδT cells relies on strong TCR signaling as well as the Signaling Lymphocyte Adaptor Molecule (SLAM)/SLAM-Associated Adaptor Protein (SAP) pathway, which induces *Zbtb16*, *Zbtb7b*, and *Klf2,* and confers upon NKγδT cells their characteristic phenotype and cytokine secretion profile ^19,33,34,35^. Vγ1.1Vδ6.3-expressing NKγδT cells expand dramatically in mice lacking downstream factors of TCR-signaling such as Itk, SLP-76, and Id3 ^19,21,36–40^. This suggests strong TCR signals emanating from the Vγ1.1Vδ6.3 TCR, likely in response to engagement by a high affinity ligand, normally lead to deletion; however, in the context of settings where TCR-signaling is attenuated by the absence of key regulators, these cells escape from negative selection and expand^8^. This interpretation was borne out by a direct test using a high affinity ligand for a defined γδ TCR, which resulted in deletion of γδ T cells that were then rescued by Id3-defiiency^8^. Nevertheless, the mechanistic basis by which Id3-defiency causes expansion of Vγ1.1Vδ6.3 NKγδT cells remains unclear.

Here we explore the mechanistic basis by which disturbing the genomic landscape by Id3-deficiency results in expansion of the Vγ1.1Vδ6.3 NKγδT cells. Indeed, we determined that the *Trav15* segment, which encodes the Vδ6.3 TCR subunit, is an E protein target whose incorporation into the Vγ1Vδ6.3 TCR is regulated by E protein binding and by generally altering the E protein activity through Id3-deficiency. Moreover, once expressed, the resulting the Vγ1Vδ6.3 TCR clonotypes that expand in Id3-deficient mice also appear to more effectively specify the innate-like NKγδT cell fate in a cell-autonomous and developmentally-unrestricted manner, providing an explanation for their expansion in Id3-deficient mice. Together, these data indicate that the ability of NKγδT cells to markedly expand in the Id3-deficient setting is entirely determined by its stereotypic Vγ1Vδ6.3 TCR complex.

## RESULTS

### Id3 loss has selective effects on Vγ subsets

TCR signal strength/duration plays a key role in separation of the αβ and γδ lineages through induction of the HLH factor Id3^4,6,8,41,42,43,44^. Interestingly, however, we, and others, have previously reported the paradoxical finding that loss of the Id3 antagonist results in the marked expansion of Vγ1Vδ6.3 NKγδT cells ^8,19,21,36^. This was not observed in E19.5 fetal thymus (Fig. S1A,B), but was quite profound in Id3-deficient adults, which exhibited a marked expansion of Vγ1^+^ γδ T cells (Fig. 1A-D). Nevertheless, development of the Vγ2^+^ T cell and skin Vγ3^+^ DETC subsets was impaired by Id3-deficiency (Fig. 1A-F). To determine if the reduction of Vγ3 DETC γδ T cells in the skin of Id3-deficient mice (Fig. 1E,F) resulted from impaired development in the thymus, we performed fetal thymic organ culture (FTOC) analysis on fetal thymic lobes from Id3^+/+^, Id3^+/-^, and Id3^-/-^ mice (Fig. 2). While the development of CD4^+^CD8^+^ (DP) αβ T lineage cells was not impaired, development of Vγ3^+^ DETC γδ T cells was impaired by Id3-deficiency (Fig. 2A,B). FTOC analysis of Id3^-/-^ thymic lobes revealed that Vγ3^+^ thymocytes were equally represented early in development (E15.5), but failed to be selected and expand over time, as indicated by the failure to induce CD122 expression (Fig. 2B-D)^43^. Moreover, Id3-deficient Vγ3^+^ progenitors exhibited a clear increase in generation of CD4^+^CD8^+^ (DP) thymocytes, suggesting that the loss of Id3 results in diversion of some cells to the αβ T cell fate (Fig. S1C,D), as we have reported previously^8^. Because the γδ TCR is downmodulated during development to the DP stage, this presumably represents an underestimate of the extent of diversion to the αβ T cell fate (Fig. S1C,D)^6,44^.

**Figure 1.**
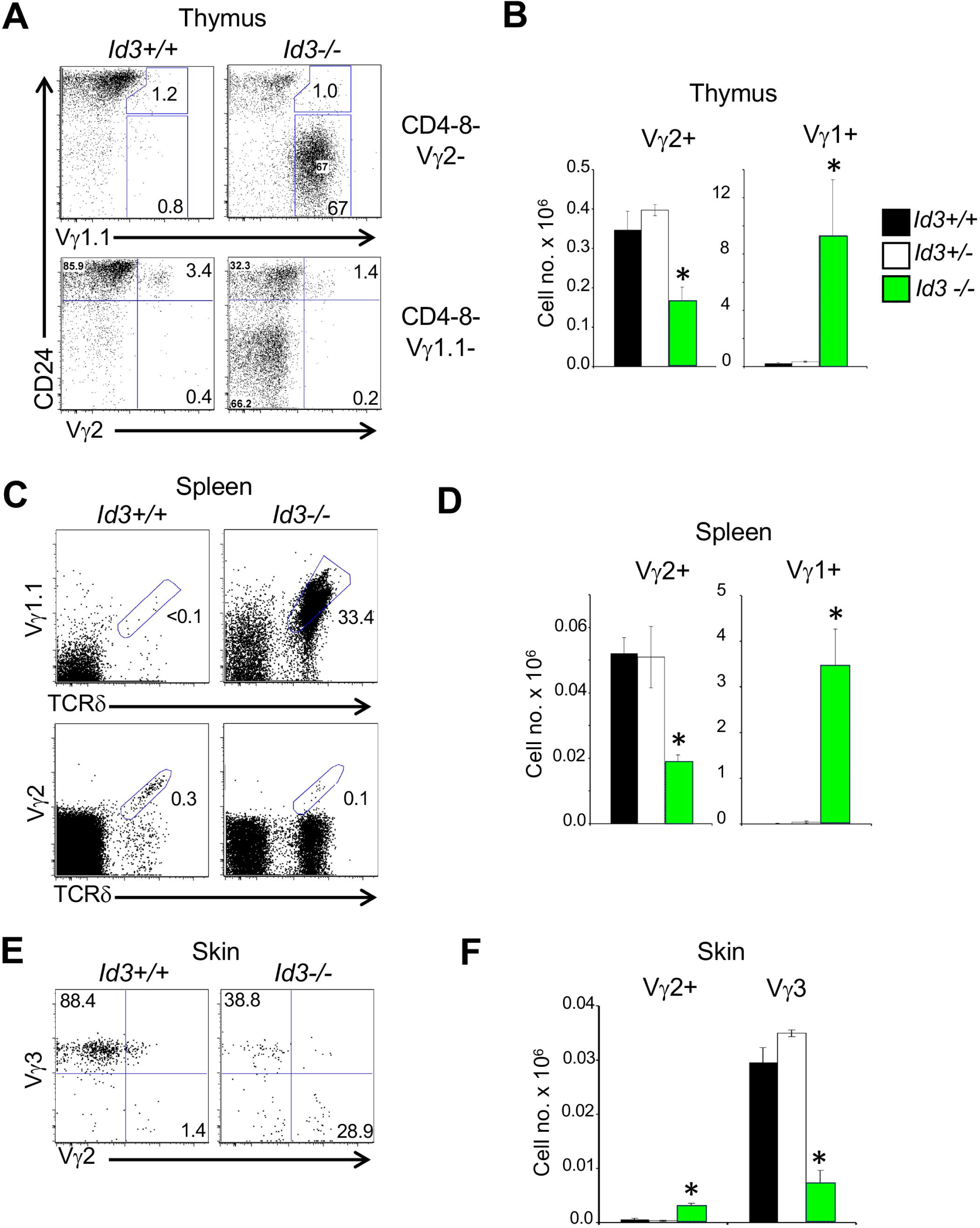
Impact of Id3-deficiency on development of γδ T cell subsets. **(A,B)** Representative flow cytometry plots of thymocytes from Id3-sufficient (Id3^+/+^) and deficient (Id3^-/-^) mice. (A) Flow cytometry plots of expression of CD24 and the indicated Vγ are displayed for CD4^-^CD8^-^Vγ2^-^ (upper panel) and CD4^-^CD8^-^Vγ1.1^-^ (lower panel) thymocytes. (B) A graphical representation is depicted of the mean absolute number ± S.D. of Vγ2^+^ and Vγ1.1^+^ thymocytes from Id3^+/+^ and Id3^-/-^ mice calculated from gate frequencies. **(C,D)** (C) Representative flow cytometry plots displaying staining for Vγ1.1 vs TCRδ, and Vγ2 vs TCRδ on splenocytes from Id3^+/+^ and Id3^-/-^ mice. (D) A graphical representation is depicted of the mean absolute number ± S.D. of Vγ2^+^ and Vγ1.1^+^ splenocytes from Id3^+/+^ and Id3^-/-^ mice calculated from gate frequencies. **(E,F)** (E) Representative flow cytometry plots of Vγ2^+^ and Vγ3^+^ staining on skin preparations from Id3^+/+^ and Id3^-/-^ mice. (F) A graphical representation is depicted of the mean absolute number ± S.D. of Vγ2^+^ and Vγ3^+^ skin γδ T cells from Id3^+/+^ and Id3^-/-^ mice calculated from gate frequencies is depicted. Data are representative of three independent experiments. Statistical significance was determined by t test. * p < 0.05

**Figure 2.**
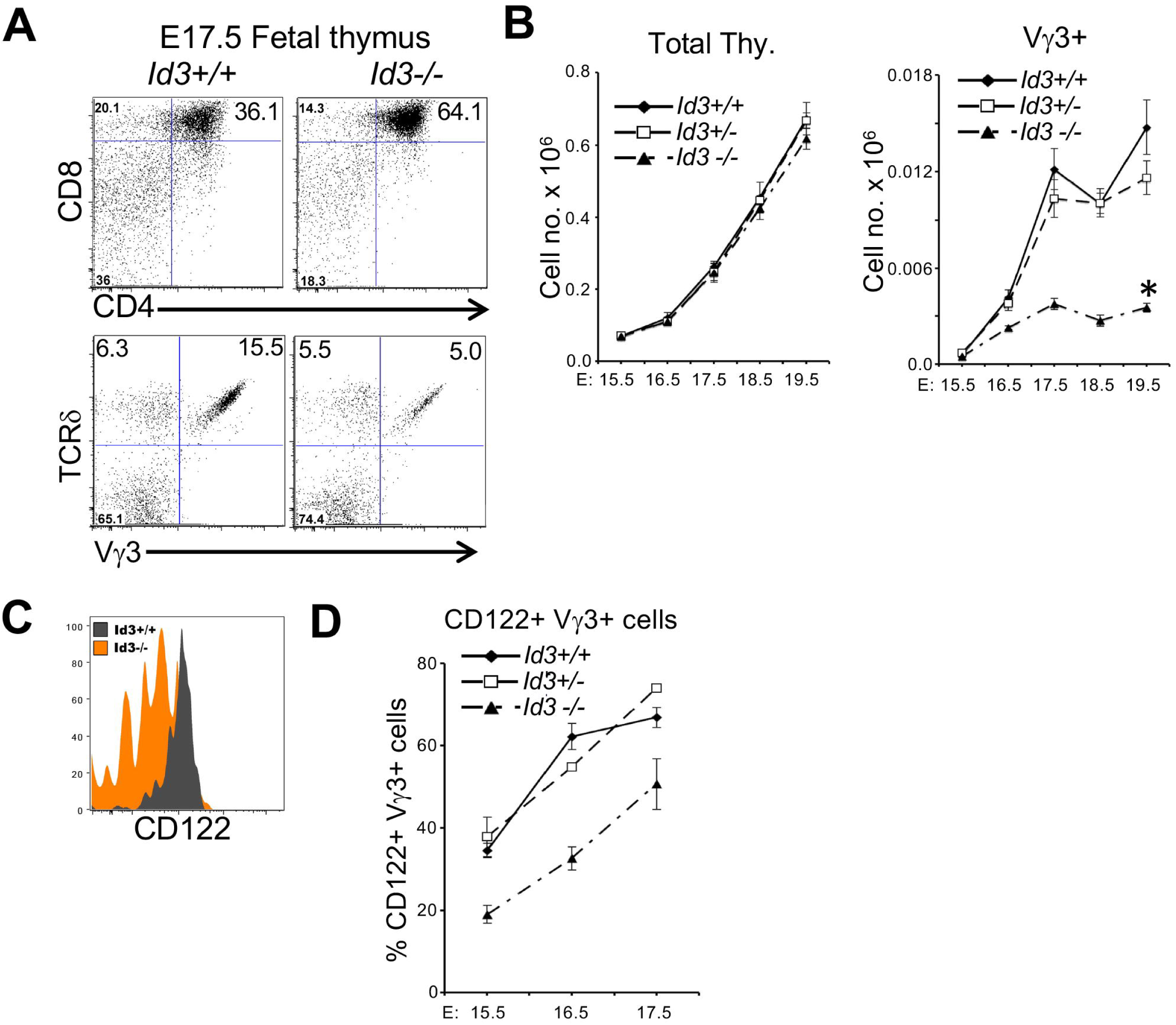
Effect of Id3-deficiency on fetal development of Vγ3^+^ γδ T cells. (A,B) (A) Representative flow cytometry plots of CD4 and CD8 staining of E17.5 thymocytes from Id3^+/+^ and Id3^-/-^ mice. (B) Graphic depiction of the mean absolute number ± S.D. of total thymocytes per lobe (left) or Vγ3^+^ γδ progenitors at E15.5 and at each of 4 additional days in fetal thymic organ culture (right). (C,D) Flow cytometry analysis of the fraction of Vγ3^+^ γδ progenitors expressing the CD122 activation marker. A representative histogram of CD122 expression is displayed (C) and mean frequency ± S.D. of the fraction of Vγ3^+^ γδ progenitors expressing CD122 is displayed graphically (D). Data are representative of three independent experiments. Statistical significance was determined by t test. * p < 0.05

### The Vγ1Vδ6.3 subset markedly expands and this is associated with CDR3 restriction

The basis for expansion of the Vγ1Vδ6.3 subset in Id3-deficient mice remains unexplained. Our previous analysis^8^ raised the possibility that a selected subclass of Vγ1Vδ6.3 TCRs, possibly with high affinity for ligand, were expanded in Id3-deficient mice. To test this possibility, we isolated immature (CD24^hi^) and mature (CD24^low^) thymic Vγ1Vδ6.3 γδ T cells from Id3^+/+^ and Id3^-/-^ mice, and sequenced their TCR γ and δ chains (Figs. 3A and S2). This revealed that the CDR3 γ and δ sequences of the Vγ1Vδ6.3 TCRs were quite diverse in immature CD24^hi^ progenitors of both Id3^+/+^ and Id3^-/-^ mice; however, upon maturation into CD24^low^ cells, the CDR3γ and δ sequences of Vγ1Vδ6.3 cells from Id3^-/-^ mice became less diverse and shorter in length, than those from Id3^+/+^ mice (Fig. 3B). The WT structures exhibit much lower pLDDT values and much greater structural variability than the KO structures. This may be because the WT sequence is longer (21 residues) than the KO sequence (17 residues), as is generally true, since WT sequences are longer and more diverse (Figure 3B). The marked restriction of the CDR3 sequences in the Vγ1Vδ6.3 TCRs of mature γδ T cells from Id3^-/-^ mice could have resulted either from a perturbation of terminal deoxynucleotide transferase (TDT) expression impacting N-region addition and/or from ligand-mediated selection of these cells.

**Figure 3.**
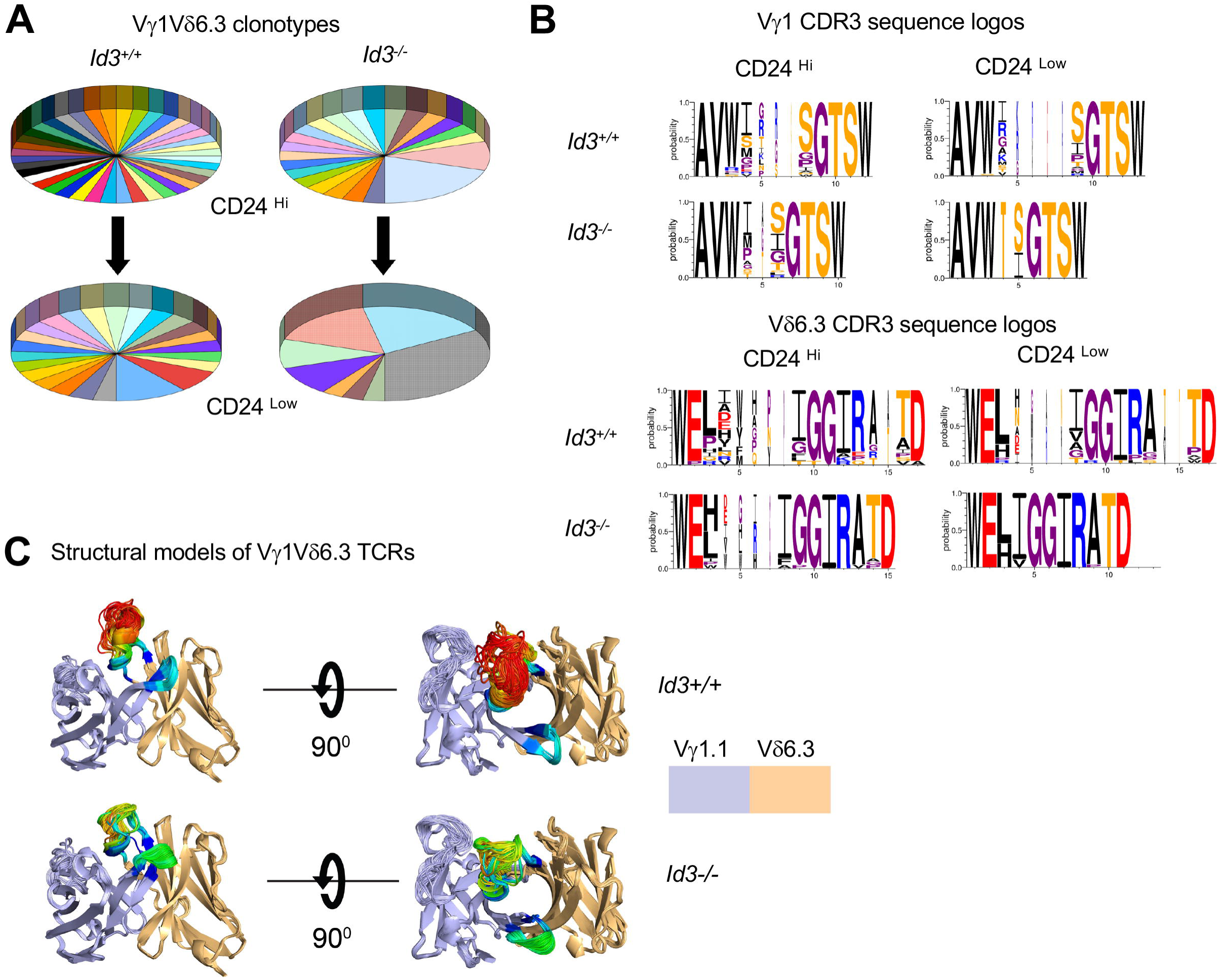
Alterations in Vγ1.1Vδ6.3 clonotypes in Id3-deficient mice. (A) Pie charts depicting the frequencies of Vγ1.1Vδ6.3 CDR3 clonotypes identified by single-cell TCR sequencing in CD24^hi^ and CD24^low^ Vγ1.1Vδ6.3 thymocytes from Id3^+/+^ and Id3^-/-^ mice. Id3^+/+^ CD24^hi^, n=40; Id3^+/+^ CD24^low^, n=37; Id3^-/-^ CD24^hi^, n=30; Id3^-/-^ CD24^low^ n=30; (B) Logos depicting the length and diversity of CDR3 sequences from the Vγ1.1 and Vδ6.3 subunits of CD24^hi^ and CD24^low^ Vγ1.1Vδ6.3 thymocytes from Id3^+/+^ and Id3^-/-^ mice. (C) Structural models of CDR3 sequences from the Vγ1.1 and Vδ6.3 subunits of CD24^low^ Vγ1.1Vδ6.3 thymocytes from Id3^+/+^ and Id3^-/-^ mice, depicting their predicted freedom of movement. Structure alignment of AlphaFold-Multimer v2.3 models of a single KO and a single WT γδ pair, chosen as representative of the sequence distribution. The frameworks and CDRs 1 and 2 are colored by TCG gene (delta=blue, gamma=orange). The CDRs are colored according to AlphaFold’s pLDDT scores, which are a measure of the predicted accuracy of the environment of each residue^71^. The coloring ranges from blue (most confident) to red (least confident).

### Vγ1Vδ6.3 TCR is sufficient to instruct the NKγδT cell fate and promote clonal dominance in Id3-deficient mice

To determine if the Vγ1Vδ6.3 TCRs that expand in Id3-deficient mice have the capacity to autonomously promote clonal dominance, we constructed inducibly-expressed Vγ1Vδ6.3 TCR transgenic (Tg) mice. In doing so, we employed one of the Vγ1Vδ6.3 TCRs that expanded among mature CD24^low^ thymocytes in Id3-deficient mice and compared it to a Vγ1Vδ6.3 TCR from Id3-sufficient mice (Fig. 4A). The Vγ1Vδ6.3 TCR Tg were knocked into the *Rosa26* locus immediately downstream of a lox-stop-lox (LSL) cassette that represses expression until Cre-mediated excision of the LSL (Fig. 4A). The TCRγ and TCRδ subunits are translated from the same transcript and the fusion protein is joined by a Tescovirus 2A (2A) sequence that enables stoichiometric expression of the separable TCRγ and TCRδ subunits (Fig. 4A). To assess the capacity of the Vγ1Vδ6.3 TCR Tg to exhibit clonal dominance, the LSL was selectively excised using 4-hydroxytamoxifen induction of *Tcrd-Cre*, which is active in a subset of T cell progenitors ^45,46^, enabling an assessment of the ability of the induced TCR to exert clonal dominance. Induction of the Vγ1Vδ6.3 TCRs derived from Id3^+/+^ (WT) and Id3^-/-^ (KO) mice in an Id3-sufficient background (*Id3^+/-^*), resulted in modest development of PLZF-expressing Vγ1Vδ6.3 NK γδ T cells, with both the WT and KO TCRs exhibiting similar ability to instruct the NKγδT cell fate (Fig. 4C-D). However, in an Id3-deficient host (*Id3^-/-^*), both TCR complexes exhibited a significantly-enhanced capacity to promote clonal dominance, as the frequency of TCRδ^+^ cells and the absolute number of cells expressing the TCR Tg, and the lineage-defining TF PLZF, was markedly increased (Fig. 4C,D). The capacity of the WT TCR to do so is somewhat surprising given its lack of predominance among the expanded clonotypes observed among mature NKγδT in Id3-deficient mice (Fig. S2). Nevertheless, consistent with its dominance in *Id3^-/-^* mice, the KO TCR exhibited a significantly greater capacity to promote adoption of the NKγδT fate (Fig. 4C,D). In parallel analysis, Id3-deficiency induced by the conditional ablation of *Id3^fl/fl^*alleles using *Tcrd-Cre^ER^*led to a less pronounced clonal expansion, presumably because of inefficient ablation of the *Id3* alleles by *Tcrd-Cre^ER^*(Fig. S3). Taken together, these data suggest that the Vγ1Vδ6.3 TCR clonotypes that preferentially expand among NKγδT cells in Id3-deficient mice are also more capable of promoting NKγδT cell development in Id3-deficient progenitors in this acute assay, even when expressed beyond the fetal/perinatal window to which their development is normally restricted^27^.

**Figure 4.**
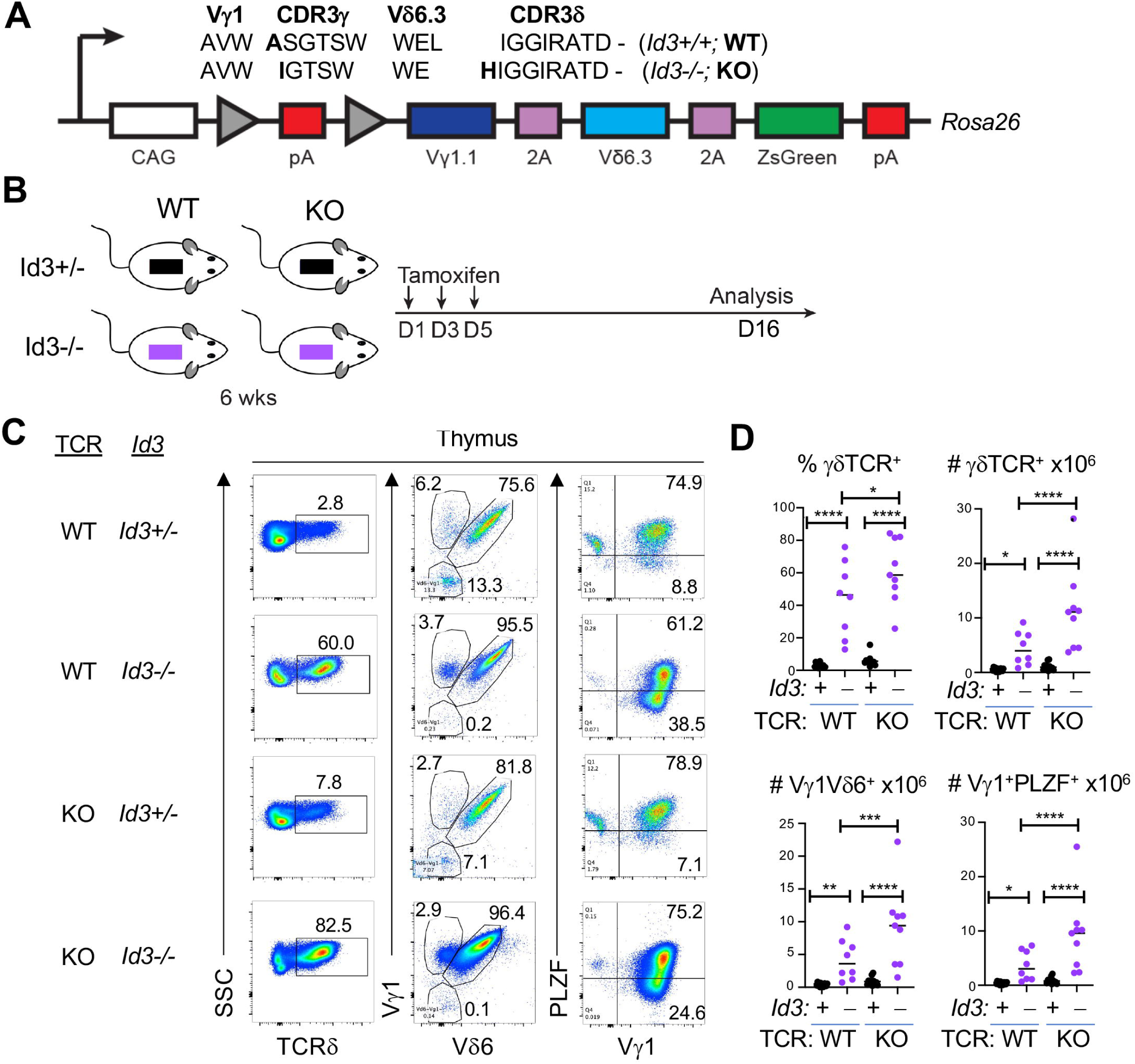
Capacity of Vγ1.1Vδ6.3 Tg from Id3^+/+^ and Id3^-/-^ mice to support development of NKγδT cells. (A) Diagram of *Rosa26* targeted Vγ1.1Vδ6.3 TCR transgene. CAG transcription is blocked by a polyadenylation (pA) sequence in the LSL element. The Vγ1.1 and Vδ6.3 TCR chains are linked by a self-cleaving Tescovirus P2A sequence. ZsGreen fluorescent reporter is linked to the TCR construct by a self-cleaving T2A sequence. The amino acid sequence comparison of the generated TCR transgenes is shown. Sequence differences are in bold. (B) Diagram of *in vivo* experimental design. Tamoxifen was administered to 6-week-old mice on days 1, 3, and 5, and analysis was performed on day 16. (C,D) (C) Representative flow cytometry plots for ZsGreen^+^ cells stained with the indicated antibodies. Frequency of γδ T lymphocytes, and absolute numbers of γδ T cells, Vγ1.1Vδ6.3 expressing cells, and Vγ1^+^PLZF^+^ γδ T cells in Id3^+/-^ and Id3^-/-^ mice are depicted graphically as scatter grams. Each dot represents an individual mouse and statistical significance was determined using two-way ANOVA with correction for multiple comparison using Tukey’s post hoc testing. WT TCR Id3^+/-^, n=14; WT TCR Id3^-/-^, n=8; KO TCR Id3^+/-^, n=11; KO TCR Id3^-/-^, n=9. *, p<0.05; **, p<0.01; ***, p<0.001; ****, p<0.0001

The KO Vγ1Vδ6.3 TCR also exhibits a greater capacity to promote the development of PLZF-expressing NKγδT cells in vitro. Cre-mediated induction of the Vγ1Vδ6.3 TCR Tg in fetal liver (FL) precursors cultured on OP9-DL1 resulted in induction of PLZF expression by progenitors expressing both the WT and KO Vγ1Vδ6.3 TCR (Fig. S4A-D). Moreover, the KO Vγ1Vδ6.3 TCR appeared to promote more extensive differentiation of mature CD73^+^CD24^low^ cells that express PLZF (Fig. S4A-D). Because NKγδT cell development is typically restricted to the late fetal/perinatal developmental window, we asked whether the KO Vγ1Vδ6.3 TCR was able to instruct development of NKγδT cells beyond the fetal/perinatal window. To test this possibility, we induced the KO Vγ1Vδ6.3 TCR in adult bone marrow progenitors and found it was able to instruct the NKγδT cell fate. Importantly, FL precursors exhibited a greater capacity to support Vγ1Vδ6.3 TCR-mediated induction of CD73 and PLZF expression than did the bone marrow progenitors (Fig. S4E-G); however, enforced expression of the Vγ1Vδ6.3 TCR in adult bone marrow progenitors was also able to promote NKγδT cell development. This suggests that once expressed, the Vγ1Vδ6.3 TCR complexes specify the NKγδT cell fate in a cell-autonomous, developmentally-unrestricted manner. Finally, while the greater permissiveness of fetal precursors to adoption of the NKγδT cell fate suggests that cellular context may influence the effectiveness with which the Vγ1Vδ6.3 TCR can instruct NKγδT cell development (Fig. S4E-G), these data indicate that restriction of NKγδT cell development to the fetal/neonatal gestational period may result from selective generation of the Vγ1Vδ6.3 TCR, perhaps under the control of the Id3/E protein axis.

### Vδ gene segment utilized to generate the Vγ1Vδ6.3 TCR

Since Vγ1^+^Vδ6.3^-^ γδ T cells continue to be generated beyond the late fetal/neonatal period, we reasoned that the restricted developmental timing of NKγδT cells may be in part regulated at the level of *Tcrd* rearrangement. NKγδT cells have been reported to have non-functional *Tcrd* rearrangements utilizing a variety of Vδ gene segments, indicating their precursors are capable of rearranging any Vδ gene segment^47^. Nevertheless, the TCRδ chain utilized by NKγδT thymocytes results almost exclusively from rearrangement of the *Trav15d-1-dv6d-1* gene segment (Vδ6.3 TCR chain), with a minority employing the *Trav15-1-dv6-1* gene segment (Vδ6.1 TCR chain) ^47^. To determine if the Id3/E axis influenced the utilization of those *Trav15* family V gene segments, E protein binding at the *Tcra-Tcrd* locus was examined by performing E2A and HEB ChIP-seq of FL derived RAG2-deficient CD4^-^CD8^-^CD44^-^CD25^+^ (DN3) T cell precursors ^48^. Interestingly, of all the Vα/Vδ segments, *Trav15d-1-dv6d-1*, *Trav15n-1*, and *Trav15-1-dv6-1* showed substantial E2A and HEB binding, which was present in both the promoter region and downstream of the RSS (Fig. 5A-C). This suggests that *Tcrd* recombination of Trav15-dv6 family V gene segments is regulated in cis by E proteins, likely by promoting chromatin accessibility. Consequently, it is possible that Id3-deficiency could influence utilization of these segments through enhancing E protein binding.

**Figure 5.**
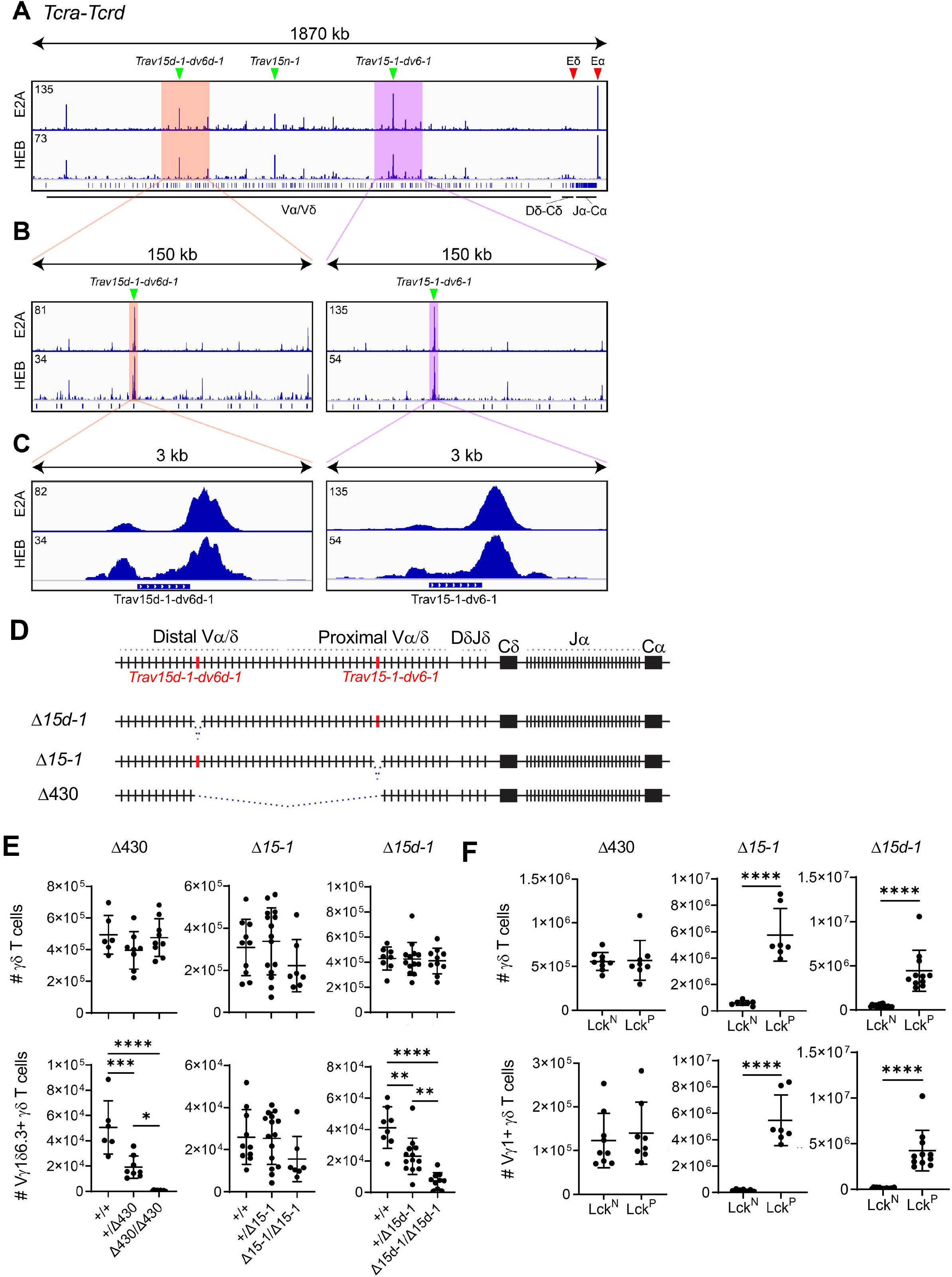
*Trav15* family members supporting development of NKγδT cells. (A-C) (A) ChIP-seq analysis of E2A and HEB binding to the C57BL/6 *Tcra-Tcrd locus* with positions of Eδ and Eα indicated by red arrows, and *Trav15d-1-dv6d-1*, *Trav15n-1*, and *Trav15-1-dv6-1* indicated by green arrowheads. (B,C) Progressively zoomed in view of the 150 kb region (B) or 3 kb region (C) highlighting the region of the *Tcra-Tcrd* locus corresponding to the *Trav15d-1-dv6d-1* and *Trav15-1-dv6-1* elements (green arrowheads). Blue bar with rightward checks indicates the location of the RSS. The maximum ChIP-seq peak value is indicated in the top left corner of the trace. Vertical blue lines in panel A denote V, D, and J gene segments, or constant region(s) as indicated. (D) Diagram of *Tcra-Tcrd* locus in 129 strain mice with relative positions of *Trav15d-1-dv6d-1* and *Trav15-1-dv6-1* indicated. Mutant alleles bearing deletions of *Trav15d-1-dv6d-1* (*Δ15d-1*), *Trav15-1-dv6-1* (*Δ15-1*), or the entire 430 kb interval (*Δ430*) are shown below. (E) Number of thymic TCRδ^+^ and Vγ1.1^+^Vδ6.3^+^ cells summarized from analysis of *Δ430, Δ15-1,* and *Δ15d-1* mutants. Littermates were used for all comparisons. *Δ15-1*: *Tcra^+/+^* (n = 10), *Tcra^+/Δ15–1^* (n = 15), and *Tcra^Δ15–1/Δ15–1^* (n = 7); *Δ15d-1: Tcra^+/+^* (n = 8), *Tcra^+/Δ15d-1^* (n = 13), and *Tcra^Δ15d-1/Δ15d-1^* (n = 10); *Δ430: Tcra^+/+^* (n = 6), *Tcra^+/Δ430^* (n = 8), and *Tcra^Δ430/Δ430^* (n = 9) Data are plotted as mean±SD and are pooled from at least 3 independent experiments. Statistical analysis: one-way ANOVA with correction for multiple comparison using Tukey’s post hoc testing. (F) The number of thymic TCRδ^+^ and Vγ1.1^+^Vδ6.3^+^ cells in Lck-Cre negative (Lck^N^) Id3 sufficient (*Id3^fl/fl^*) (control), or Lck-Cre (Lck^P^) mediated Id3 deficient (*Id3^fl/fl^*) *Δ430, Δ15-1*, and *Δ15d-1* mutants. All comparisons to Lck-Cre negative (Lck^N^) littermates. For *Δ430*: Lck^N^ (n = 9), Lck^P^ (n = 8); *Δ15-1*: Lck^N^ (n = 7), Lck^P^ (n = 7); *Δ15d-1:* Lck^N^ (n = 16) and Lck^P^ (n = 11). Data are pooled from at least 3 independent experiments and plotted as mean±SD. Statistical analysis: Student’s t test. \*\**p* < 0.01, \*\*\**p* < 0.001, \*\*\*\**p* < 0.0001.

To identify the *Trav15* element required to support development of NKγδT cells, we targeted them for deletion by CRISPR/Cas9 in *Tcrd^CreER^*mice, which lack the *Trav15n-1* segment^45^. The individual deletions of *Trav15d-1-dv6d-1* (Δ15d-1) and *Trav15-1-dv6-1* (Δ15-1) spanned ∼2.1 kb over the respective gene segment and eliminated the E protein binding regions both in the promoter and downstream of the RSS (Figures 5D). We additionally produced a *Tcra-Tcrd* allele with a 430 kb deletion within the Vα/Vδ array (*Δ430*), which lacks both of the regions deleted in the *Δ15d-1* and *Δ15-1* mutants, as well as the intervening 430 kb with the V gene segments contained therein (Figure 5D). None of the resulting mutant mice exhibited alterations in thymic cellularity, CD4/8 subsets, or total γδ T cells (Figs. 5E and S5A,B). As expected, ablation of both Trav15-1 family gene segments (Δ*430*) blocked development of NKγδT cells (Fig. 5E). Importantly, while ablation of the proximal *Trav15* segment, *Trav15-1-dv6-1* (*Δ15-1*), failed to block development of NKγδT cells, their development was severely attenuated by ablation of the distal segment, *Trav15d-1-dv6d-1* (*Δ15d-1*) (Fig. 5E), demonstrating that NKγδT cell development depends predominantly on use of *Trav15d-1-dv6d-1* (Fig. 5E).

We next sought to determine whether globally-enhancing E protein activity through Id3-deficiency would alter the capacity of *Trav15* family members to support expansion of γδ T cells expressing the Vγ1Vδ6.3 TCR. Id3-deficiency in Tcra^Δ430/Δ430^ mice did not lead to observable differences in thymic cellularity or γδ T lymphocytes (Figs. 5F and S5C); however, elimination of Id3 in Tcra^Δ15–1/Δ15–1^ mice resulted in a significant reduction of thymic cellularity and a significant expansion of Vγ1.1^+^ T lymphocytes (Figs. 5F and S5C), consistent with our finding that NKγδT cell development depends primarily on *Trav15d-1-dv6d-1*, which is intact in these mice. Interestingly, Id3-deletion in Tcra^Δ15d–1/Δ15d-1^ also resulted in a significant reduction of thymic cellularity and a significant expansion of Vγ1.1^+^ T lymphocytes which were largely PLZF^+^ (Fig. S5D) and thus appear to be NKγδT cells whose expansion was supported by the Vγ1.1Vδ6.1 TCR. Taken together, these data indicate that development of NKγδT is primarily dependent upon the *Trav15d-1-dv6d-1* element for encoding the TCRδ subunit of their stereotyped Vγ1.1Vδ6.3 TCR complex; however, when E protein activity is enhanced by Id3-deficiency and the *Trav15d-1-dv6d-1* element is eliminated, then the *Trav15-1-dv6-1* element is capable of supporting NKγδT cell development through a Vγ1.1Vδ6.1 TCR complex.

### E protein bound elements supporting NKγδT cell development

Having determined that generally enhancing E protein activity altered the usage of *Trav15* gene segments by developing NKγδT cells, we wished to assess the role of cis-acting E protein binding sties adjacent to the *Trav15* element (*Trav15d-1-dv6d-1)* preferentially utilized by Vγ1.1Vδ6.3 expressing NKγδT cells. There are three consensus E box binding sites within the first 100 bp downstream of the RSS of the *Trav15d-1-dv6d-1* gene segment (Fig. 6A). To assess their role in supporting the *Trav15d-1-dv6d-1-*encoded Vδ6.3 subunit of the TCR that supports NKγδT cell development, the E-box containing region downstream of *Trav15d-1-dv6d-1* was ablated by CRISPR editing (Fig. 6A; Δ*E*). It should be noted that to eliminate the possibility of compensation by the *Trav15-1-dv6-1* element, we ablated the E protein binding sites on the *Δ15-1* allele described above. The Δ*E* mutation did not affect the total number of γδ T cells (Fig. 6B); however, it did reduce both the frequency and number of Vγ1Vδ6.3 expressing NKγδT cells (Fig. 6B), strongly suggesting that E protein binding downstream of *Trav15d-1-dv6d-1* element plays a critical role in regulating recombination at this *Trav15* element and supporting the expression of the Vγ1Vδ6.3 TCR preferred by NKγδT cells. Because predicted binding sites for other TFs mapped to the 59 bp ablated in Δ*E*, such as Sox4 and Runx1, we directly tested the role of the E protein binding sites by selectively inactivating the 1^st^ (Δ*E1*) and 3^rd^ (Δ*E3* and *mE3*) E box sites (Fig. 6B). As with deletion of the entire 59 bp interval, inactivation of the 1^st^ or 3^rd^ E box sites did not alter the total number of γδ T cells (Fig. 6B); however, inactivation of those E box sites did reduce both the frequency and number of Vγ1Vδ6.3 expressing NKγδT cells (Fig. 6B).

**Figure 6.**
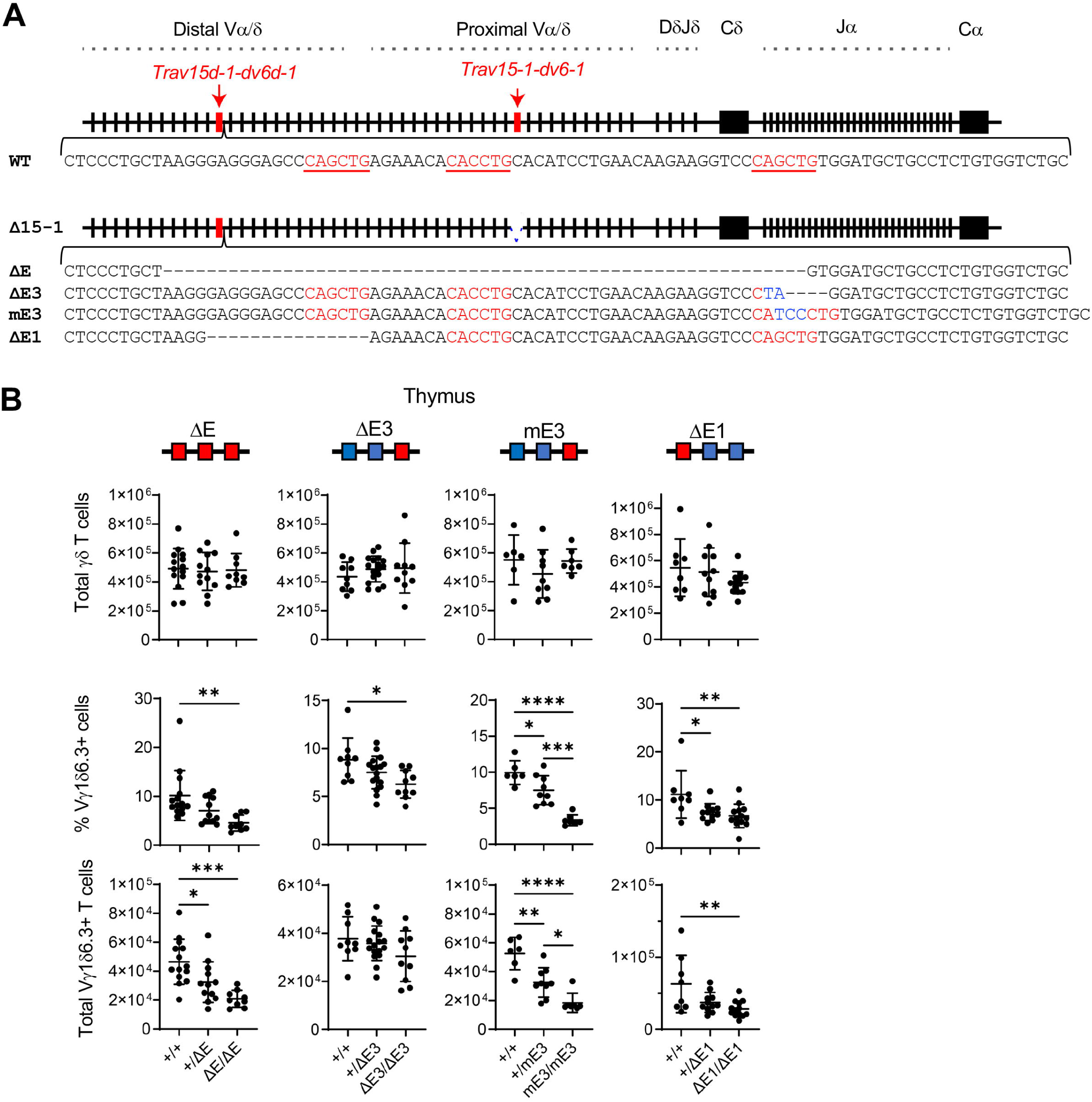
Role of E protein binding in selection of *Trav15* family V genes during NKγδT cell development. (A) Diagram of *Tcra-Tcrd* locus in 129 strain mice with relative positions of *Trav15d-1-dv6d-1* and *Trav15-1-dv6-1* indicated. The region containing E-boxes adjacent to *Trav15d-1-dv6d-1* RSS was mutated on the *Δ15-1* allele to prevent compensatory usage of the *Trav15-1-dv6-1* element. Mutations to the Ebox binding sites are indicated. (B) The number of thymic TCRδ^+^, and frequency and number of thymic Vγ1.1^+^Vδ6.3^+^ cells for each of the E box mutant mice (*ΔE, ΔE3, mE3, and ΔE1*) are depicted graphically as scatter plots. All comparisons represent littermates. *ΔE: Tcra^+/+^* (n = 14), *Tcra^+/ΔE^*(n = 12), and *Tcra^ΔE/ΔE^*(n = 9); *ΔE3: Tcra^+/+^*(n = 8), *Tcra^+/ΔE3^*(n = 17), and *Tcra^ΔE3/ΔE3^*(n = 10); mE3: *Tcra^+/+^*(n = 6), *Tcra^+/mE3^*(n = 9), and *Tcra^mE3/mE3^*(n = 7); *ΔE1: Tcra^+/+^*(n = 8), *Tcra^+/ΔE1^*(n = 11), and *Tcra^ΔE1/ΔE1^*(n = 13). Data were pooled from at least 3 independent experiments and are plotted as mean±SD. Statistical analysis: one-way ANOVA with correction for multiple comparison using Tukey’s post hoc testing. \**p* < 0.05, \*\**p* < 0.01, \*\*\**p* < 0.001, \*\*\*\**p* < 0.0001.

Taken together, our analysis indicates the Id3/E protein axis plays an important role in both generation of the Vγ1Vδ6.3 TCR complex and its capacity to specify the NKγδT cell fate.

## DISCUSSION

Previous analysis strongly supports the hypothesis that TCR signals of differing strength regulate the separation of the αβ and γδ lineage fates through Id3-mediated graded repression of the function of E proteins, with weak signals causing modest Id3 induction, retention of significant E protein activity, and adoption of the αβ fate while strong TCR signals markedly repress E protein activity and promote the γδ fate^8,48,49^. While the central role of Id3 in αβ/γδ lineage commitment is supported by studies with defined γδ TCR complexes^8^ and for the Vγ3^+^ DETC cells whose development is attenuated by Id3-deficiency, development of Vγ1Vδ6.3 expressing NKγδT cell stands in stark contrast, as NKγδT cells are not only not dependent on Id3, but are actually restrained by it^8,19–21^. Here we provide an explanation for this conundrum, revealing that the Id3/E protein axis plays a key role in regulating the development of NKγδT cells, by controlling both the generation of the stereotyped NKγδT Vγ1Vδ6.3 TCR complex, and its capacity to support NKγδT cell development.

Our sequence analysis revealed that the Vγ1.1Vδ6.3 TCR complexes of the NKγδT cells that expanded in Id3-deficient mice were characterized by CDR3 sequences that were shorter and less diverse than those found in Id3-sufficient mice. One explanation for this may be related to the absence of TDT, which is required for non-templated addition of nucleotides (N) to the CDR3 junctions^50,51^. The Vγ1.1Vδ6.3 TCR complexes of NKγδT cells may lack these N-additions either because TDT is not expressed during fetal gestation when NKγδT cells develop^50^, or because E proteins regulate TDT expression, as has been reported in B lineage cells^52^. Importantly, E protein regulation of TDT seems unlikely to explain our observation of shorter CDR3 elements, since Id3-deficiency enhances E protein activity, which has been reported to increase TDT expression^52^ and should thus increase N-region additions and CDR3 length. Accordingly, a more likely explanation for the expansion in Id3-deficient mice of NKγδT cells with restricted CDR3s is a change in competitive fitness of progenitors bearing Vγ1.1Vδ6.3 TCR complexes in the context of Id3-deficiency, perhaps in response to ligand-mediated selection. We previously tested this possibility using the KN6 γδ TCR Tg mice, showing that development of γδ T cells in response to TCR signals induced by low affinity ligands was blocked by Id3-deficiency, while development in response to TCR signals elicited by high-affinity ligands that normally delete or restrain γδ T cell development in Id3-sufficient mice, promoted expansion in Id3-deficient mice^8^. Likewise, we determined that conditional expression of Vγ1.1Vδ6.3 TCR complexes in adult Id3-sufficient mice (6wk old) was able to instruct the development of PLZF-expressing NKγδT cells, albeit in limited numbers. Importantly, when the Vγ1.1Vδ6.3 TCR complexes were induced in Id3-deficient mice, development of PLZF-expressing NKγδT cells was markedly enhanced. Both the Vγ1.1Vδ6.3 TCR complexes isolated from Id3-sufficient and Id3-deficient mice were able to promote NKγδT cell development, although the Vγ1.1Vδ6.3 TCR complexes isolated from Id3-defiicient (KO) mice exhibited a significantly greater ability to do so, providing an explanation for its preferential expansion in Id3-deficient mice.

The ability of the Vγ1.1Vδ6.3 TCR complexes to support the profound expansion of NKγδT cells even in the context of Id3-deficiency raises the question of why they support expansion when other γδ TCR complexes (KN6 and Vγ3^+^ DETC) are unable to do so^8^. The restricted length and diversity of the CDR3 sequences of the TCRγ and TCRδ chains likely stems from a TCR-ligand mediated selection, as has been previously suggested^8,47^. The putative intrathymic ligand for Vγ1.1Vδ6.3 TCR complexes has not been identified, but it likely involves polyanionic features, as polyanions have been shown to activate Vγ1.1Vδ6.3 TCRs in vitro^53^. The Vγ1.1Vδ6.1 and Vγ1.1Vδ6.3 TCRs likely recognize the same ligand, with the latter possibly having higher binding affinity which could support their ability to outcompete Vγ1.1Vδ6.1 T lymphocytes for intrathymic encounters with a high-affinity ligand. In this scenario, Itk- or Id3-deficiency attenuate the capacity of a strong TCR-ligand interaction to transmit a signal capable of promoting developmental arrest or death^8,40^. We have recently showed that the Id3-deficiency does so through effects on E proteins that involve a regulatory loop controlling the induction of Egr2 and Myc, which are required for expansion of NKγδT cells in the absence of Id3^54–56^. Together, these data indicate that Id3-deficiency influences the capacity of Vγ1.1Vδ6.3 TCR complexes to support the profound expansion of NKγδT cells.

Our finding that enforced expression of Vγ1.1Vδ6.3 TCR complexes instructs the profound expansion of NKγδT cells, even when expressed outside of the fetal/perinatal window to which their generation is normally restricted, strongly suggests that the developmental restriction of NKγδT cell generation is controlled by the developmental timing of expression of the Vγ1.1Vδ6.3 TCR. Consequently, we also explored the basis for usage of the *Trav15* family V genes utilized to generate the Vδ6.3 subunit, since the Vγ1.1 usage is not restricted to the fetal/perinatal development. Individual deletions of the *Trav15d-1-dv6d-1* and *Trav15-1-dv6-1* gene segments supported previous sequencing-based investigation suggesting that the Vδ6.3 subunit of the NKγδT cell TCR is encoded by the *Trav15d-1-dv6d-1* gene segment^31,47^. Importantly, the *Trav15-1-dv6-1* segment could substitute for the *Trav15d-1-dv6d-1* gene in supporting NKγδT cell development if E protein activity was globally augmented through Id3-deficiency and the *Trav15d-1-dv6d-1* gene segment is absent. This suggested that E protein activity was controlling *Trav15* family rearrangement, and could be occurring by local control, since we found strong E2A and HEB binding peaks near the *Trav15* elements. We found that deletion or disruption of the E-box(es) downstream of the *Trav15d-1-dv6d-1* RSS impaired Vγ1.1Vδ6.3 T lymphocyte development indicating that E proteins are regulating *Trav15* V gene element utilization directly through cis-acting regulatory elements, but this does not provide an explanation for the strong preference for the *Trav15d-1-dv6d-1* element. There are two potential mechanisms by which E protein action at the *Trav15* family elements could support the preferential utilization of *Trav15d-1-dv6d-1* in supporting the development of Vγ1.1Vδ6.3 NKγδT cells. First, E proteins could selectively promote V(D)J recombination of particular *Trav15* family members through preferential binding; however, our E protein ChIP-seq analysis revealed strong E protein binding to the RSS sequences adjacent to both *Trav15d-1-dv6d-1* and *Trav15-1-dv6-1,* which is inconsistent with this notion. Moreover, our deletion analysis clearly showed that the disfavored *Trav15-1-dv6-1* gene segment could only support development of NKγδT cells if both the preferred *Trav15d-1-dv6d-1* gene segment and Id3 were ablated. Finally, NKγδT thymocytes have been reported to contain non-functional *Tcrd* rearrangements utilizing a variety of Vδ gene segments, indicating their precursors are capable of rearranging any Vδ gene segment, but that only particular Vδ gene segments can support NKγδT development ^47^. The second possibility is that the TCRδ subunit resulting from rearrangement of *Trav15d-1-dv6d-1* confers a competitive advantage to the resulting Vγ1.1Vδ6.3 TCR expressing progenitors. Interestingly, the CDR1, CDR2 and adjacent sequences encoded by the *Trav15d-1-dv6d-1* and *Trav15-1-dv6-1* elements differ by 10 amino acids (aa), including several charged aa and a bulky proline residue. These charged residues might impact the extent of interaction with polyanionic surfaces, which have been implicated in activation of Vγ1.1Vδ6.3 TCRs^53^. Accordingly, it is possible that the *Trav15d-1-dv6d-1* and *Trav15-1-dv6-1* elements are rearranged with the same frequency, but that the ten aa that differ between the Vδ6.3 and Vδ6.1 subunits confer a selective advantage, perhaps in response to ligand engagement. Similarly, germline encoded motifs in mouse and human Vγ subunits have been implicated in promoting γδ T cell activation^57,58^, and development through interaction with butyrophilin-like ligands^18,59–61^. The importance of these 10 aa differences in development and the putative ligand with which they might interact is under investigation.

The signal strength model stipulates that the induction of Id3 in proportion to TCR signal strength results in graded reductions in E protein activity that play a critical role in separation of the αβ and γδ lineages. While there is significant data in support of this model and critical role played by Id3 in supporting these fate decisions, Vγ1.1Vδ6.3 expressing NKγδT cells represent an exception since their development does not require Id3. On the contrary, their development is restrained by Id3. We provide here clear evidence that Id3 plays a clear role in both generating the stereotyped Vγ1.1Vδ6.3 TCR that promotes NKγδT cell development and its capacity to drive the marked expansion of NKγδT cells in the context of Id3-deficiency. Importantly, the Vγ1.1Vδ6.3 TCR can instruct the NKγδT fate in a cell autonomous manner at any developmental window where it is expressed, indicating the developmental restriction of this lineage is regulated at the level of Vγ1.1Vδ6.3 TCR generation. Greater insight into the relative competitive fitness of particular Vγ1.1Vδ6.3 TCR complexes in supporting NKγδT development in the absence of Id3 will ultimately depend on identifying the putative selecting ligand(s).

## ACKNOWLEGEMENTS

Finally, we gratefully acknowledge the assistance of the following core facilities of the Fox Chase Cancer Center: Cell Culture, Flow Cytometry, Laboratory Animal, and Molecular Modeling. We also gratefully acknowledge the assistance of the Flow Cytometry Shared Resource and the Transgenic and Knockout Mouse Shared Resource of the Duke University Cancer Institute, and the Duke University Division of Laboratory Animal Resources. DLW was supported by NIH grant P01AI102853, core grant P30CA006927, the Bishop Fund and an appropriation from the Commonwealth of Pennsylvania. RLD was supported by R35GM122517, and YZ was supported by NIH grants P01AI102853 and R01GM059638.

## AUTHOR CONTRIBUTIONS

DLW, YZ, and MC conceived and oversaw the study and together with AM wrote the manuscript. AM, SYL, SS, AVC, BZ, MR, MIP and RD performed experiments and analyzed data. JCZP contributed critical reagents.

## MATERIALS AND METHODS

### Mice

Generation of the *Tcrd^CreER^*, *Id3^-/-^*, *Id3^fl/fl^*, and Lck^Cre^ lines was described previously ^45,62–64^. All mice were maintained in AALAC-accredited laboratory animal facilities at either Fox Chase Cancer Center or Duke University. All experiments were conducted under institutional animal care and use committee (IACUC) approved protocols.

### Tissue isolation and flow cytometry

Flow cytometry was performed on single cell suspensions from thymus, spleen and epidermis as described^65^. Cells were isolated and stained with the following antibodies: anti-CD3 (17A2), anti-CD90.2 (30-H12), anti-CD4 (GK1.5 or RM4-5), anti-CD8 (53-6.7), anti-CD24 (M1/69), anti-CD73 (TY/11.8), anti-CD122 (TM-b1), anti-TCRβ (H57-597), anti-TCRδ (GL3), anti-Vγ1 (2.11), anti-Vγ2 (UC3-10A6), anti-Vγ3 (536), anti-Vd6.3 (C504.17C), anti-PLZF (9E12), anti-NK1.1 (PK136), anti-IL4 (11B11) and anti-IFNγ (XMG1.2). The antibodies were purchased from eBioscience, BD Biosciences, or BioLegend. Conventional flow cytometry was performed as described using DAPI or propidium iodide (PI) to exclude dead cells^48^. Intracellular flow cytometry for cytokine production was performed as previously described^48^. Cytokine production was assessed by intracellular staining of splenocytes using anti-IL4 and anti-IFNγ antibodies after stimulation for 30 minutes with 50 ng/ml Phorbol 12-myristate 13-acetate (PMA) and 1 μg/ml Ionomycin followed by culture for another 5 hours and 30min with 1 μg Brefeldin A. Data were analyzed on either an LSRII, Fortessa X20 or Symphony A5 flow cytometer (BD Biosciences). FlowJo Software (TreeStar) was used to analyze data. Cell populations were purified by flow cytometry using a FACSAria Cell Sorter (BD Biosciences).

### Fetal thymic organ culture (FTOC)

Following timed mating of mice, fetal thymic lobes were isolated from mice at E15.5, following which they were cultured *in vitro* at the air-medium interface, as described^6^. Developmental progression was monitored daily by flow cytometry on triplicate thymic lobes.

### Single cell TCR sequencing

Individual CD24^hi^ or CD24^low^ Vγ1.1^+^ γδ T cells were isolated from *Id3^+/+^ and Id3^-/-^* mice by flow cytometry and deposited in wells of glass slides (Beckman-Coulter). Vγ1.1 and Vδ6.3 subunits were then amplified by RT-PCR using nested primers in single cells: Vγ1 external F-gggcttgggcagctggagca; Cγ4 R-gaaggaaggaaaatagtagg; Vγ1 internal F-agtatctaatatatgtctca; Cγ4 R-ggagaaaagtctgagtcagt; Vδ6 external F-ggatctaatgtggccgaga; Cδ R-tgttccatttttcatgatga; Vδ6 internal F-aagtgattcaggtctggtca; Cδ R-tggtttggccggaggctggc. Following amplification, the cDNA fragments were sequenced to assess the impact of Id3-deficiency on the CDR3 sequences. Following isolation, the Vγ1.1 and Vδ6.3 pairs were cloned into pMiCherry as Tescovirus 2A-linked Vγ1.1Vδ6.3 fusion proteins, as described^8^.

### Structural modeling of the Vγ1.1Vδ6.3 TCR complexes

Structure predictions were performed with AlphaFold-Multimer v2.3 (https://www.biorxiv.org/content/10.1101/2021.10.04.463034v2.abstract) as implemented in ColabFold running on local machines^66^. The structures of each γδ. TCR were predicted with 20 random seeds without templates with all five AlphaFold-Multimer models, for a total of 100 predictions with each. Models were relaxed with AMBER within ColabFold. Models with pTM scores lower than 0.8 were excluded, since they involved unphysiological domain-swapped dimers and other anomalies. Figures were created within PyMol. Sequence logos were produced with WebLogo^67^. Sequences for each group (KO vs WT x CD24^hi^ vs CD24^low^) were aligned manually in a text editor. Sequences were aligned to the longest sequences such that the length of the sequence logo is the length of the longest sequence. Identical sequences were not pruned before producing the sequence logo.

### Generation of mutant mouse lines

A Vγ1.1Vδ6.3 TCR complex that exhibited expanded representation among CD24^low^ mature Vγ1.1Vδ6.3 expressing cells from Id3-deficient mice (KO TCR) and a Vγ1.1Vδ6.3 complex from Id3-sufficient mice that was not expanded in Id3-deficient mice (WT TCR), were selected for production of inducible TCR Tg mice. The WT and KO 2A-linked Vγ1.1Vδ6.3 TCR constructs that were preceded by a lox-stop-lox (LSL) sequence were cloned between the short and long-arm of the *Rosa26* locus in the Ai6 targeting vector. G4 mouse embryonic stem (ES) cells were then transfected with the Ai6 targeting vector to knock the TCR constructs into the *Rosa26* locus. The targeted ES cells were injected into blastocytes by the Duke University Cancer Institute Transgenic and Knockout Mouse Shared Resource and the resulting chimeric mice were screened for transmission of the targeted allele to offspring. The *Trav15d-1-dv6d-1* and *Trav15-1-dv6-1* gene segment deletions, and the *Tcra-Tcrd* locus 430 kb deletion (*Tcra^Δ4^*^30^ allele) were generated using the same CRISPR/Cas9 targeting strategy reported previously^68^. Briefly, the deletions were generated using two gRNA flanking the element to be targeted (5′-TCTTCCCTTAAAGAGTGATA-3′ and 5′-GACATTAGAGTCCCTTAAAG-3′) by CRISPR/Cas9 electroporation into embryos from *Tcrd^CreER/CreER^;Id3^fl/fl^;R26^ZsG/ZsG^*mice on a C57BL/6 strain background^45,63,69^. Founders were screened by Sanger sequencing and maintained separately by crossing to *Tcrd^CreER/CreER^;Id3^fl/fl^;R26^ZsG/ZsG^*. Deletions of the E-box containing region downstream of *Trav15d-1-dv6d-1* were similarly generated in *Tcra^Δ15–1/Δ15–1^;Id3^fl/fl^;R26^ZsG/ZsG^*mice using two-guides flanking the target sequences to be ablated (5′-TGCAGGTGTGTTTCTCAGCT-3′ and 5′-GAACAAGAAGGTCCCAGCTG-3′). Founders were screened by Sanger sequencing and maintained separately by crossing to *Tcrd^CreER/CreER^;Id3^fl/fl^;R26^ZsG/ZsG^*.

### Tamoxifen induction of the Vγ1.1Vδ6.3 TCR Tg

WT and KO Vγ1.1Vδ6.3 Tg on Id3-sufficient or Id3-deficient backgrounds were induced by tamoxifen treatment in vivo. Mice were injected i.p. with 1mg of tamoxifen on days 1, 3, and 5, following which support of NKγδT cell development was assessed by flow cytometry on either day 16 or 17, as indicated. TCR Tg induction was monitored using a Zs-Green LSL reporter cassette. For induction of Vγ1.1Vδ6.3 Tg *in vitro*, either E16.5 FL or adult bone marrow were isolated from Vγ1.1Vδ6.3 Tg mice. FLs were disaggregated by manual disruption. Lineage negative hematopoietic stem and progenitor cells (HSPC) were isolated from bone marrow using magnetic bead depletion with biotinylated antibodies (B220, CD3, CD4, CD8, CD11b, CD11c, CD19, CD44, GR-1, IgM, NK1.1, TCRβ, and TCRγδ) and streptavidin magnetic beads. To evaluate the capacity of the WT and KO Vγ1.1Vδ6.3 Tg to promote adoption of the NKγδT fate *in vitro*, FL or bone marrow HSPC were cultured on OP9-DL1 cells in the presence of 1 ng/mL IL-7 and 5 ng/mL FLT3L (BioLegend) as previously described ^70^. After 8 days in polarizing conditions, cells were placed in experimental conditions. Tg TCRs were induced using 4-hydroxytamoxifen (250 ng/mL). Cells were cultured on OP9-DL1 cells in the presence of 1 ng/mL IL-7 and 5 ng/mL FLT3L until day 6 post-induction, and only in the presence of 1 ng/mL IL-7 thereafter (BioLegend). Cultures were analyzed and passaged every 3 days.

### ChIP-seq analysis

Chip-Seq for E2A and HEB was performed on DN3 cell generated from *Rag2^-/-^* FL progenitors cultured on OP9-DL1 monolayers^48^. The resulting ChIP-seq analysis was performed as described and has been deposited in GEO (GEO: GSE162292)^48^. The resulting data were reanalyzed here. Alignment was performed using Bowtie2, peak calling was performed using MACS2, following which data were visualized using Integrative Genomics Viewer.

### Statistical analysis

Sample size was not predetermined by statistical methods. Statistical significance was determined using GraphPad Prism, which was also used for generation of graphs.

---

**Figure S1. Impact of Id3-deficiency on fetal development of Vγ1.1^+^ γδ T cells**

(A,B) (A) Representative flow cytometry plots depicting the frequency of Vγ1.1^+^ γδ T cells in E19.5 fetal thymus. thymocytes for *Id3^+/+^, Id3^+/-^,* and *Id3^-/-^*mice. (B) The mean ± SD of the absolute number of Vγ1.1^+^ γδ T cells per lobe at E18.5 and E19.5 is depicted graphically.

(C,D) Diversion of Vγ3^+^ thymocytes γδ T to the αβ fate was assessed by gating on Vγ3^+^ expressing thymocytes (upper panel) and displaying CD4 and CD8 expression (lower panel).

The mean ± SD of the frequency of Vγ3^+^ cells diverted to the αβ T cell fate, as indicated by becoming CD4^+^8^+^ double positive thymocytes, is depicted graphically for E17.5, E18.5, and E19.5 thymocytes for mice of the indicated genotypes.

Representative flow cytometry plots are displayed. All data represent at least 3 independent experiments. Statistical significance was assessed by t-test. \*\**p* < 0.01.

**Figure S2. Effect of Id3 deficiency on the diversity of CDR3γ and CDR3δ diversity of the Vγ1.1Vδ6.3 TCRs.**

Single cell sequencing was performed for the V regions of the Vγ1.1 and Vδ6.3 subunits of CD24^hi^ immature and CD24^low^ mature γδ T cells from *Id3^+/+^* and *Id3^-/-^* mice. The sequences of the clonotypes are listed relative to germline sequence, and their representation relative to the total number of clonotypes sequences is indicated in parenthesis on the right. The clonotype sequence of the Vγ1.1Vδ6.3 TCRs used to make the WT and KO Tg mice are highlighted in yellow.

**Figure S3. Impact of conditional ablation of *Id3* using TCRd-Cre on development of Vγ1.1Vδ6.3 TCR expressing NKγδT cells.**

(A) Diagram of *in vivo* experimental design. Tamoxifen was administered to 6-week-old mice on days 1, 3, and 5, and analysis was performed on day 17.

(B) Graphic depiction of the frequency of ZsGreen^+^ thymocytes.

(C) Representative flow cytometry plots of ZsGreen^+^TCRδ^+^ thymocytes displayed as Vγ1.1 vs Vδ6.3 for mice expressing the WT or KO Vγ1.1Vδ6.3 TCR.

(D,E) Number of thymic (D) or splenic (E) TCRδ^+^, Vγ1.1^+^Vδ6.3^+^, and PLZF^+^ Vγ1.1^+^Vδ6.3^+^ cells is depicted graphically with each symbol representing a single mouse.

(F) Graphic representation of the frequency and number of IL-4 and IFNγ co-producing Vγ1.1^+^Vδ6.3^+^ splenocytes following PMA/Ionomycin stimulation, with each symbol representing a single mouse.

*TCR^WT^Id3^+/+^*(n = 10), *TCR^WT^Id3^fl/fl^*(n = 7), *TCR^KO^Id3^+/+^*(n = 9), and *TCR^KO^Id3^fl/fl^*(n = 9). Data were pooled from 3 independent experiments and are plotted as mean±SD. Statistical analysis: two-way ANOVA with correction for multiple comparison using Tukey’s post hoc testing \**p* < 0.05, \*\**p* < 0.01, \*\*\**p* < 0.001, \*\*\*\**p* < 0.0001.

**Figure S4. Capacity of Vγ1.1Vδ6.3 TCR complexes to instruct the NKγδT cell fate in vitro.**

(A) Representative flow cytometry plots of FL derived Vγ1.1Vδ6.3 T lymphocyte maturation in culture displayed as CD73 vs CD24. The WT and KO Tg TCRs were induced in FL derived precursors cultured in T cell polarizing conditions.

(B-D) Graphic depiction of the (B) absolute numbers Vγ1.1Vδ6.3 expressing cells obtained, (C) frequency which were CD73^+^CD24^low^, and (D) frequency of Vγ1.1Vδ6.3 cells expressing PLZF.

(E-G) The KO Tg TCR was induced in FL and bone marrow derived precursors cultured in T cell polarizing conditions. Graphic depiction of the (E) absolute numbers Vγ1.1Vδ6.3 expressing cells obtained, and the frequencies of these which were (F) CD73^+^CD24^low^ or (G) expressed PLZF.

Data are pooled from 3 independent experiments (n = 6). Data are plotted as mean±SD. Statistical analysis: two-way ANOVA with correction for multiple comparison using Sidak’s multiple comparisons test. \**p* < 0.05, \*\**p* < 0.01, \*\*\**p* < 0.001, \*\*\*\**p* < 0.0001.

**Figure S5. Role of E protein binding in *Trav15* element selection in developing NKγδT cells.**

(A) Thymic cellularity in *Δ430, Δ15-1*, and *Δ15d-1* mutant mice is depicted graphically with each symbol representing an individual mouse. All comparisons involved littermates. *Δ15-1: Tcra^+/+^* (n = 10), *Tcra^+/Δ15–1^* (n = 15), and *Tcra^Δ15-1/Δ15-1^* (n = 7); *Δ15d-1: Tcra^+/+^*(n = 8), *Tcra^+/Δ15d-1^* (n = 13), and *Tcra^Δ15d-1/Δ15d-1^* (n = 10); *Δ430: Tcra^+/+^* (n = 6), *Tcra^+/Δ430^* (n = 8), and *Tcra^Δ430/Δ430^*(n = 9). Data were pooled from a minimum of 3 independent experiments and are plotted as mean±SD. Statistical analysis: one-way ANOVA with correction for multiple comparison using Tukey’s post hoc testing. Significant differences were not detected.

(B) Representative flow cytometry plots for subsets defined by CD4 and CD8 expression in *Tcra*^+/+^, *Tcra^Δ430/Δ430^*, *Tcra^Δ15-1/Δ15-1^, Tcra^Δ15d-1/Δ15d-1^* mutant mice.

(C) Thymic cellularity in Lck-Cre negative (Lck^N^) Id3 sufficient (*Id3^fl/fl^*) (control), or Lck-Cre (Lck^P^) mediated Id3 deficient (*Id3^fl/fl^*) *Δ430, Δ15-1,* and *Δ15d-1* mutants. All comparisons to Lck-Cre negative (Lck^N^) littermates. For *Δ430:* Lck^N^ (n = 9), Lck^P^ (n = 8); *Δ15-1*: Lck^N^ (n = 7), Lck^P^ (n = 7); *Δ15d-1:* Lck^N^ (n = 16) and Lck^P^ (n = 11). Data are pooled from at least 3 independent experiments and plotted as mean±SD. Statistical analysis: Student’s t test. \*\*\*\**p* < 0.0001.

(D) Representative flow cytometry plots of NK1.1 and PLZF expression in Vγ1^+^ thymocytes from Lck-Cre negative (Lck^N^) Id3 sufficient (*Id3^fl/fl^*) (control), or Lck-Cre (Lck^P^) mediated Id3 deficient (*Id3^fl/fl^*) *Δ430, Δ15-1,* and *Δ15d-1* mutants

(E) Thymic cellularity in *ΔE, ΔE3, mE3,* and *ΔE1* mutant mice is depicted graphically with each symbol representing an individual mouse. All comparisons involved littermates. *ΔE: Tcra^+/+^*(n = 14), *Tcra^+/ΔE^*(n = 12), and *Tcra^ΔE/ΔE^*(n = 9); *ΔE3: Tcra^+/+^*(n = 8), *Tcra^+/ΔE3^*(n = 17), and *Tcra^ΔE3/ΔE3^* (n = 10); *mE3: Tcra^+/+^* (n = 6), *Tcra^+/mE3^* (n = 9), and *Tcra^mE3/mE3^* (n = 7); *ΔE1: Tcra^+/+^* (n = 8), *Tcra^+/ΔE1^* (n = 11), and *Tcra^ΔE1/ΔE1^*(n = 13). Data were pooled from a minimum of 3 independent experiments and are plotted as mean±SD. Statistical analysis: one-way ANOVA with correction for multiple comparison using Tukey’s post hoc testing. Significant differences were not detected.

## LITERATURE CITED

1. Parker, M.E., and Ciofani, M. (2020). Regulation of γδ T Cell Effector Diversification in the Thymus. Front Immunol 11, 42. 10.3389/fimmu.2020.00042.

2. Ciofani, M., and Zúñiga-Pflücker, J.C. (2010). Determining γδ versus αß T cell development. Nat Rev Immunol 10, 657–663. 10.1038/nri2820.

3. Ciofani, M., Knowles, G.C., Wiest, D.L., von Boehmer, H., and Zúñiga-Pflücker, J.C. (2006). Stage-specific and differential notch dependency at the alphabeta and gammadelta T lineage bifurcation. Immunity 25, 105–116. 10.1016/j.immuni.2006.05.010.

4. Hayes, S.M., Li, L., and Love, P.E. (2005). TCR signal strength influences alphabeta/gammadelta lineage fate. Immunity 22, 583–593. 10.1016/j.immuni.2005.03.014.

5. Lee, S.Y., Stadanlick, J., Kappes, D.J., and Wiest, D.L. (2010). Towards a molecular understanding of the differential signals regulating alphabeta/gammadelta T lineage choice. Semin Immunol. 22, 237–246. doi: 210.1016/j.smim.2010.1004. 1008. Epub 2010 May 1014.

6. Haks, M.C., Lefebvre, J.M., Lauritsen, J.P., Carleton, M., Rhodes, M., Miyazaki, T., Kappes, D.J., and Wiest, D.L. (2005). Attenuation of gammadeltaTCR signaling efficiently diverts thymocytes to the alphabeta lineage. Immunity 22, 595–606. 10.1016/j.immuni.2005.04.003.

7. Lee, S.Y., Coffey, F., Fahl, S.P., Peri, S., Rhodes, M., Cai, K.Q., Carleton, M., Hedrick, S.M., Fehling, H.J., Zúñiga-Pflücker, J.C., et al. (2014). Noncanonical mode of ERK action controls alternative αβ and γδ T cell lineage fates. Immunity. 41, 934–946. doi: 910.1016/j.immuni.2014.1010. 1021. Epub 2014 Nov 1028.

8. Lauritsen, J.P., Wong, G.W., Lee, S.Y., Lefebvre, J.M., Ciofani, M., Rhodes, M., Kappes, D.J., Zúñiga-Pflücker, J.C., and Wiest, D.L. (2009). Marked induction of the helix-loop-helix protein Id3 promotes the gammadelta T cell fate and renders their functional maturation Notch independent. Immunity 31, 565–575. 10.1016/j.immuni.2009.07.010.

9. Murre, C. (2019). Helix-loop-helix proteins and the advent of cellular diversity: 30 years of discovery. Genes Dev 33, 6–25. 10.1101/gad.320663.118.

10. Anderson, M.K., and da Rocha, J.D.B. (2022). Direct regulation of TCR rearrangement and expression by E proteins during early T cell development. WIREs Mech Dis 14, e1578. 10.1002/wsbm.1578.

11. Miyazaki, M., and Miyazaki, K. (2022). The E-Id axis specifies adaptive and innate lymphoid lineage cell fates. J Biochem 172, 259–264. 10.1093/jb/mvac068.

12. Miyazaki, K., Watanabe, H., Yoshikawa, G., Chen, K., Hidaka, R., Aitani, Y., Osawa, K., Takeda, R., Ochi, Y., Tani-Ichi, S., et al. (2020). The transcription factor E2A activates multiple enhancers that drive Rag expression in developing T and B cells. Sci Immunol 5. 10.1126/sciimmunol.abb1455.

13. Kadakia, T., Tai, X., Kruhlak, M., Wisniewski, J., Hwang, I.Y., Roy, S., Guinter, T.I., Alag, A., Kehrl, J.H., Zhuang, Y., and Singer, A. (2019). E-protein-regulated expression of CXCR4 adheres preselection thymocytes to the thymic cortex. J Exp Med 216, 1749–1761. 10.1084/jem.20182285.

14. In, T.S.H., Trotman-Grant, A., Fahl, S., Chen, E.L.Y., Zarin, P., Moore, A.J., Wiest, D.L., Zúñiga-Pflücker, J.C., and Anderson, M.K. (2017). HEB is required for the specification of fetal IL-17-producing γδ T cells. Nat Commun 8, 2004 10.1038/s41467-017-02225-5.

15. Miyazaki, M., Miyazaki, K., Chen, K., Jin, Y., Turner, J., Moore, A.J., Saito, R., Yoshida, K., Ogawa, S., Rodewald, H.R., et al. (2017). The E-Id Protein Axis Specifies Adaptive Lymphoid Cell Identity and Suppresses Thymic Innate Lymphoid Cell Development. Immunity 46, 818–834.e814. 10.1016/j.immuni.2017.04.022.

16. Belle, I., and Zhuang, Y. (2014). E proteins in lymphocyte development and lymphoid diseases. Curr Top Dev Biol 110, 153–187. 10.1016/b978-0-12-405943-6.00004-x.

17. Zhang, B., Lin, Y.Y., Dai, M., and Zhuang, Y. (2014). Id3 and Id2 act as a dual safety mechanism in regulating the development and population size of innate-like γδ T cells. J Immunol 192, 1055–1063. 10.4049/jimmunol.1302694.

18. Contreras, A.V., and Wiest, D.L. (2020). Recent advances in understanding the development and function of γδ T cells. 1000Res. 9:F1000, 10.12688/f11000research.22161.12681. eCollection 12020.

19. Alonzo, E.S., Gottschalk, R.A., Das, J., Egawa, T., Hobbs, R.M., Pandolfi, P.P., Pereira, P., Nichols, K.E., Koretzky, G.A., Jordan, M.S., and Sant’Angelo, D.B. (2010). Development of promyelocytic zinc finger and ThPOK-expressing innate gamma delta T cells is controlled by strength of TCR signaling and Id3. J Immunol 184, 1268–1279. 10.4049/jimmunol.0903218.

20. Verykokakis, M., Boos, M.D., Bendelac, A., and Kee, B.L. (2010). SAP protein-dependent natural killer T-like cells regulate the development of CD8(+) T cells with innate lymphocyte characteristics. Immunity 33, 203–215. 10.1016/j.immuni.2010.07.013.

21. Ueda-Hayakawa, I., Mahlios, J., and Zhuang, Y. (2009). Id3 restricts the developmental potential of gamma delta lineage during thymopoiesis. J Immunol 182, 5306–5316. 10.4049/jimmunol.0804249.

22. Havran, W.L., and Allison, J.P. (1988). Developmentally ordered appearance of thymocytes expressing different T-cell antigen receptors. Nature 335, 443–445. 10.1038/335443a0.

23. Ito, K., Bonneville, M., Takagaki, Y., Nakanishi, N., Kanagawa, O., Krecko, E.G., and Tonegawa, S. (1989). Different gamma delta T-cell receptors are expressed on thymocytes at different stages of development. Proc Natl Acad Sci U S A 86, 631–635. 10.1073/pnas.86.2.631.

24. Ribot, J.C., Lopes, N., and Silva-Santos, B. (2021). γδ T cells in tissue physiology and surveillance. Nat Rev Immunol 21, 221–232. 10.1038/s41577-020-00452-4.

25. Havran, W.L., Chien, Y.H., and Allison, J.P. (1991). Recognition of self antigens by skin-derived T cells with invariant gamma delta antigen receptors. Science 252, 1430–1432. 10.1126/science.1828619.

26. Itohara, S., Farr, A.G., Lafaille, J.J., Bonneville, M., Takagaki, Y., Haas, W., and Tonegawa, S. (1990). Homing of a gamma delta thymocyte subset with homogeneous T-cell receptors to mucosal epithelia. Nature 343, 754–757. 10.1038/343754a0.

27. Contreras, A.V., and Wiest, D.L. (2023). Development of γδ T Cells: Soldiers on the Front Lines of Immune Battles. Methods Mol Biol 2580:71-88., 10.1007/1978-1001-0716-2740-1002_1004.

28. Lees, R.K., Ferrero, I., and MacDonald, H.R. (2001). Tissue-specific segregation of TCRgamma delta+ NKT cells according to phenotype TCR repertoire and activation status: parallels with TCR alphabeta+NKT cells. Eur J Immunol 31, 2901–2909. 10.1002/1521-4141(2001010)31:10<2901::aid-immu2901>3.0.co;2-#.

29. Vicari, A.P., Mocci, S., Openshaw, P., O’Garra, A., and Zlotnik, A. (1996). Mouse gamma delta TCR+NK1.1+ thymocytes specifically produce interleukin-4, are major histocompatibility complex class I independent, and are developmentally related to alpha beta TCR+NK1.1+ thymocytes. Eur J Immunol 26, 1424–1429. 10.1002/eji.1830260704.

30. Gerber, D.J., Azuara, V., Levraud, J.P., Huang, S.Y., Lembezat, M.P., and Pereira, P. (1999). IL-4-producing gamma delta T cells that express a very restricted TCR repertoire are preferentially localized in liver and spleen. J Immunol 163, 3076–3082.

31. Azuara, V., Levraud, J.P., Lembezat, M.P., and Pereira, P. (1997). A novel subset of adult gamma delta thymocytes that secretes a distinct pattern of cytokines and expresses a very restricted T cell receptor repertoire. Eur J Immunol 27, 544–553. 10.1002/eji.1830270228.

32. Grigoriadou, K., Boucontet, L., and Pereira, P. (2003). Most IL-4-producing gamma delta thymocytes of adult mice originate from fetal precursors. J Immunol 171, 2413–2420. 10.4049/jimmunol.171.5.2413.

33. Kreslavsky, T., Savage, A.K., Hobbs, R., Gounari, F., Bronson, R., Pereira, P., Pandolfi, P.P., Bendelac, A., and von Boehmer, H. (2009). TCR-inducible PLZF transcription factor required for innate phenotype of a subset of gammadelta T cells with restricted TCR diversity. Proc Natl Acad Sci U S A 106, 12453–12458. 10.1073/pnas.0903895106.

34. Dienz, O., DeVault, V.L., Musial, S.C., Mistri, S.K., Mei, L., Baraev, A., Dragon, J.A., Krementsov, D., Veillette, A., and Boyson, J.E. (2020). Critical Role for SLAM/SAP Signaling in the Thymic Developmental Programming of IL-17- and IFN-γ-Producing γδ T Cells. J Immunol 204, 1521–1534. 10.4049/jimmunol.1901082.

35. Anderson, M.K., and Selvaratnam, J.S. (2020). Interaction between γδTCR signaling and the E protein-Id axis in γδ T cell development. Immunol Rev 298, 181–197. 10.1111/imr.12924.

36. Verykokakis, M., Boos, M.D., Bendelac, A., Adams, E.J., Pereira, P., and Kee, B.L. (2010). Inhibitor of DNA binding 3 limits development of murine slam-associated adaptor protein-dependent "innate" gammadelta T cells. PLoS One 5, e9303. 10.1371/journal.pone.0009303.

37. Jordan, M.S., Smith, J.E., Burns, J.C., Austin, J.E., Nichols, K.E., Aschenbrenner, A.C., and Koretzky, G.A. (2008). Complementation in trans of altered thymocyte development in mice expressing mutant forms of the adaptor molecule SLP76. Immunity 28, 359–369. 10.1016/j.immuni.2008.01.010.

38. Qi, Q., Xia, M., Hu, J., Hicks, E., Iyer, A., Xiong, N., and August, A. (2009). Enhanced development of CD4+ gammadelta T cells in the absence of Itk results in elevated IgE production. Blood 114, 564–571. 10.1182/blood-2008-12-196345.

39. Felices, M., Yin, C.C., Kosaka, Y., Kang, J., and Berg, L.J. (2009). Tec kinase Itk in gammadeltaT cells is pivotal for controlling IgE production in vivo. Proc Natl Acad Sci U S A 106, 8308–8313. 10.1073/pnas.0808459106.

40. Yin, C.C., Cho, O.H., Sylvia, K.E., Narayan, K., Prince, A.L., Evans, J.W., Kang, J., and Berg, L.J. (2013). The Tec kinase ITK regulates thymic expansion, emigration, and maturation of γδ NKT cells. J Immunol 190, 2659–2669. 10.4049/jimmunol.1202531.

41. Zarin, P., Wong, G.W., Mohtashami, M., Wiest, D.L., and Zúñiga-Pflücker, J.C. (2014). Enforcement of γδ-lineage commitment by the pre-T-cell receptor in precursors with weak γδ-TCR signals. Proc Natl Acad Sci U S A 111, 5658–5663. 10.1073/pnas.1312872111.

42. Garbe, A.I., Krueger, A., Gounari, F., Zúñiga-Pflücker, J.C., and von Boehmer, H. (2006). Differential synergy of Notch and T cell receptor signaling determines alphabeta versus gammadelta lineage fate. J Exp Med 203, 1579–1590. 10.1084/jem.20060474.

43. Xiong, N., Kang, C., and Raulet, D.H. (2004). Positive selection of dendritic epidermal gammadelta T cell precursors in the fetal thymus determines expression of skin-homing receptors. Immunity 21, 121–131. 10.1016/j.immuni.2004.06.008.

44. Ferrero, I., Mancini, S.J., Grosjean, F., Wilson, A., Otten, L., and MacDonald, H.R. (2006). TCRgamma silencing during alphabeta T cell development depends upon pre-TCR-induced proliferation. J Immunol 177, 6038–6043. 10.4049/jimmunol.177.9.6038.

45. Zhang, B., Wu, J., Jiao, Y., Bock, C., Dai, M., Chen, B., Chao, N., Zhang, W., and Zhuang, Y. (2015). Differential Requirements of TCR Signaling in Homeostatic Maintenance and Function of Dendritic Epidermal T Cells. J Immunol 195, 4282–4291. 10.4049/jimmunol.1501220.

46. Zhang, B., Jia, Q., Bock, C., Chen, G., Yu, H., Ni, Q., Wan, Y., Li, Q., and Zhuang, Y. (2016). Glimpse of natural selection of long-lived T-cell clones in healthy life. Proc Natl Acad Sci U S A 113, 9858–9863. 10.1073/pnas.1601634113.

47. Azuara, V., Lembezat, M.P., and Pereira, P. (1998). The homogeneity of the TCRdelta repertoire expressed by the Thy-1dull gammadelta T cell population is due to cellular selection. Eur J Immunol 28, 3456–3467. 10.1002/(sici)1521-4141(199811)28:11<3456::Aid-immu3456>3.0.Co;2-f.

48. Fahl, S.P., Contreras, A.V., Verma, A., Qiu, X., Harly, C., Radtke, F., Zúñiga-Pflücker, J.C., Murre, C., Xue, H.H., Sen, J.M., and Wiest, D.L. (2021). The E protein-TCF1 axis controls γδ T cell development and effector fate. Cell Rep 34, 108716. 10.1016/j.celrep.2021.108716.

49. Fahl, S.P., Kappes, D.J., and Wiest, D.L. (2018). TCR Signaling Circuits in αβ/γδ T Lineage Choice.

50. Bogue, M., Gilfillan, S., Benoist, C., and Mathis, D. (1992). Regulation of N-region diversity in antigen receptors through thymocyte differentiation and thymus ontogeny. Proc Natl Acad Sci U S A. 89, 11011–11015. doi: 11010.11073/pnas.11089.11022.11011.

51. Komori, T., Okada, A., Stewart, V., and Alt, F.W. (1993). Lack of N regions in antigen receptor variable region genes of TdT-deficient lymphocytes. Science. 261, 1171–1175. doi: 1110.1126/science.8356451.

52. Choi, J.K., Shen, C.P., Radomska, H.S., Eckhardt, L.A., and Kadesch, T. (1996). E47 activates the Ig-heavy chain and TdT loci in non-B cells. Embo j 15, 5014–5021.

53. Dunst, J., Glaros, V., Englmaier, L., Sandoz, P.A., Önfelt, B., Kisielow, J., and Kreslavsky, T. (2020). Recognition of synthetic polyanionic ligands underlies "spontaneous" reactivity of Vγ1 γδTCRs. J Leukoc Biol. 107, 1033–1044. doi: 1010.1002/JLB.1032MA1219-1392R. Epub 2020 Jan 1013.

54. Roy, S., Moore, A.J., Love, C., Reddy, A., Rajagopalan, D., Dave, S.S., Li, L., Murre, C., and Zhuang, Y. (2018). Id Proteins Suppress E2A-Driven Invariant Natural Killer T Cell Development prior to TCR Selection. Front Immunol 9, 42. 10.3389/fimmu.2018.00042.

55. Zhang, B., Jiao, A., Dai, M., Wiest, D.L., and Zhuang, Y. (2018). Id3 Restricts γδ NKT Cell Expansion by Controlling Egr2 and c-Myc Activity. J Immunol 201, 1452–1459. 10.4049/jimmunol.1800106.

56. Li, J., Wu, D., Jiang, N., and Zhuang, Y. (2013). Combined deletion of Id2 and Id3 genes reveals multiple roles for E proteins in invariant NKT cell development and expansion. J Immunol 191, 5052–5064. 10.4049/jimmunol.1301252.

57. Karunakaran, M.M., Willcox, C.R., Salim, M., Paletta, D., Fichtner, A.S., Noll, A., Starick, L., Nöhren, A., Begley, C.R., Berwick, K.A., et al. (2020). Butyrophilin-2A1 Directly Binds Germline-Encoded Regions of the Vγ9Vδ2 TCR and Is Essential for Phosphoantigen Sensing. Immunity 52, 487–498.e486. 10.1016/j.immuni.2020.02.014.

58. Willcox, C.R., Vantourout, P., Salim, M., Zlatareva, I., Melandri, D., Zanardo, L., George, R., Kjaer, S., Jeeves, M., Mohammed, F., et al. (2019). Butyrophilin-like 3 Directly Binds a Human Vγ4(+) T Cell Receptor Using a Modality Distinct from Clonally-Restricted Antigen. Immunity 51, 813–825.e814. 10.1016/j.immuni.2019.09.006.

59. Di Marco Barros, R., Roberts, N.A., Dart, R.J., Vantourout, P., Jandke, A., Nussbaumer, O., Deban, L., Cipolat, S., Hart, R., Iannitto, M.L., et al. (2016). Epithelia Use Butyrophilin-like Molecules to Shape Organ-Specific γδ T Cell Compartments. Cell 167, 203–218.e217. 10.1016/j.cell.2016.08.030.

60. Barbee, S.D., Woodward, M.J., Turchinovich, G., Mention, J.J., Lewis, J.M., Boyden, L.M., Lifton, R.P., Tigelaar, R., and Hayday, A.C. (2011). Skint-1 is a highly specific, unique selecting component for epidermal T cells. Proc Natl Acad Sci U S A 108, 3330–3335. 10.1073/pnas.1010890108.

61. Boyden, L.M., Lewis, J.M., Barbee, S.D., Bas, A., Girardi, M., Hayday, A.C., Tigelaar, R.E., and Lifton, R.P. (2008). Skint1, the prototype of a newly identified immunoglobulin superfamily gene cluster, positively selects epidermal gammadelta T cells. Nat Genet 40, 656–662. 10.1038/ng.108.

62. Pan, L., Hanrahan, J., Li, J., Hale, L.P., and Zhuang, Y. (2002). An analysis of T cell intrinsic roles of E2A by conditional gene disruption in the thymus. J Immunol 168, 3923–3932. 10.4049/jimmunol.168.8.3923.

63. Guo, Z., Li, H., Han, M., Xu, T., Wu, X., and Zhuang, Y. (2011). Modeling Sjögren’s syndrome with Id3 conditional knockout mice. Immunol Lett 135, 34–42. 10.1016/j.imlet.2010.09.009.

64. Pan, L., Sato, S., Frederick, J.P., Sun, X.H., and Zhuang, Y. (1999). Impaired immune responses and B-cell proliferation in mice lacking the Id3 gene. Mol Cell Biol. 19, 5969–5980. doi: 5910.1128/MCB.5919.5969.5969.

65. Chen, E.L.Y., Lee, C.R., Thompson, P.K., Wiest, D.L., Anderson, M.K., and Zúñiga-Pflücker, J.C. (2021). Ontogenic timing, T cell receptor signal strength, and Notch signaling direct γδ T cell functional differentiation in vivo. Cell Rep 35, 109227. 10.1016/j.celrep.2021.109227.

66. Mirdita, M., Schütze, K., Moriwaki, Y., Heo, L., Ovchinnikov, S., and Steinegger, M. (2022). ColabFold: making protein folding accessible to all. Nat Methods. 19, 679–682. doi: 610.1038/s41592-41022-01488-41591. Epub 42022 May 41530.

67. Crooks, G.E., Hon, G., Chandonia, J.M., and Brenner, S.E. (2004). WebLogo: a sequence logo generator. Genome Res. 14, 1188–1190. doi: 1110.1101/gr.849004.

68. Dauphars, D.J., Mihai, A., Wang, L., Zhuang, Y., and Krangel, M.S. (2022). Trav15-dv6 family Tcrd rearrangements diversify the Tcra repertoire. J Exp Med 219. 10.1084/jem.20211581.

69. Madisen, L., Zwingman, T.A., Sunkin, S.M., Oh, S.W., Zariwala, H.A., Gu, H., Ng, L.L., Palmiter, R.D., Hawrylycz, M.J., Jones, A.R., et al. (2010). A robust and high-throughput Cre reporting and characterization system for the whole mouse brain. Nat Neurosci 13, 133–140. 10.1038/nn.2467.

70. Holmes, R., and Zúñiga-Pflücker, J.C. (2009). The OP9-DL1 system: generation of T-lymphocytes from embryonic or hematopoietic stem cells in vitro. Cold Spring Harb Protoc 2009, pdb.prot5156. 10.1101/pdb.prot5156.

71. Mariani, V., Biasini, M., Barbato, A., and Schwede, T. (2013). lDDT: a local superposition-free score for comparing protein structures and models using distance difference tests. Bioinformatics. 29, 2722–2728. doi: 2710.1093/bioinformatics/btt2473. Epub 2013 Aug 2727.

