## Supplementary data for "E protein control of NKγδT cell development through both generation and function of the stereotypic Vγ1Vδ6.3 TCR"

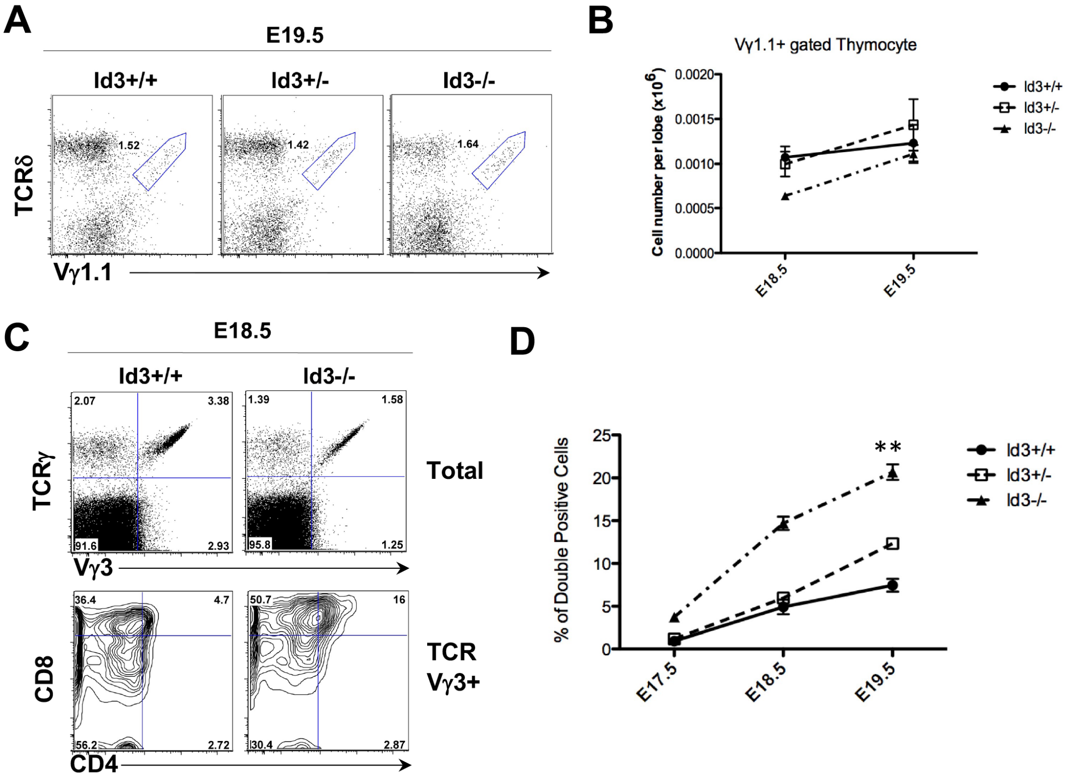


**Figure S1. Impact of Id3-deficiency on fetal development of Vγ1.1$^+$ γδ T cells**

(A,B) (A) Representative flow cytometry plots depicting the frequency of Vγ1.1$^+$ γδ T cells in E19.5 fetal thymus. thymocytes for *Id3$^{+/+}$, Id3$^{+/-}$,* and *Id3$^{-/-}$* mice. (B) The mean ± SD of the absolute number of Vγ1.1$^+$ γδ T cells per lobe at E18.5 and E19.5 is depicted graphically.

(C,D) Diversion of Vγ3$^+$ thymocytes γδ T to the αβ fate was assessed by gating on Vγ3$^+$ expressing thymocytes (upper panel) and displaying CD4 and CD8 expression (lower panel). The mean ± SD of the frequency of Vγ3$^+$ cells diverted to the αβ T cell fate, as indicated by becoming CD4$^+$8$^+$ double positive thymocytes, is depicted graphically for E17.5, E18.5, and E19.5 thymocytes for mice of the indicated genotypes.

Representative flow cytometry plots are displayed. All data represent at least 3 independent experiments. Statistical significance was assessed by t-test. ***p* < 0.01.

ID3 WT (CD24Hi) Singlecel PCR

| Vγ1.1 | P/N | Jγ4 | Vδ6.3 | P/N | Dδ1 | P/N | Dδ2 | P/N | Jδ1 | |
|---|---|---|---|---|---|---|---|---|---|---|
| GCA GTC TGG ATA AA | | TCA GGC ACA TCA TGG | TGG GAG CTG G | | ATGGCATAT | | ATCGGAGGGATACGAG | | CT ACC GAC | Gemline |
| GCA GTC TGG | TCA | TCA GGC ACA TCA TGG | TGG GAG CTG G | | | TTAC | CGGAGGGATACGAG | CAACGG | CC GAC | |
| GCA GTC TGG ATA A | GA | TCA GGC ACA TCA TGG | TGG GAG CT | | | | CGGAGG | CCGACAAA | CT ACC GAC | |
| GCA GTC TGG ATA | GGA | TCA GGC ACA TCA TGG | TGG GAG C | | | CGCG AT | ATCGGAGGG | CCG | CT ACC GAC | |
| GCA GTC TGG | | TCA GGC ACA TCA TGG | TGG GAG C | GAATGA | GGCATAT | ATTAC | GGG | ACG | CT ACC GAC | |
| GCA GTC TGG ATA AA | GGGA | TCA GGC ACA TCA TGG | TGG GAG CT | CATA | ATGGC | CCCCAT | ATCGGAGGGATACG | CCC | CT ACC GAC | |
| GCA GTC TGG | TCTCGA | TCA GGC ACA TCA TGG | TGG GAG CTG G | TA | GG | AG | ATCGGAGGGATACGA | CAC | CT ACC GAC | |
| GCA GTC TGG AT | CCCC | TCA GGC ACA TCA TGG | TGG GAG | ATC | GCAT | TTT | TCGGAGGGATACGA | AGAATTG | CT ACC GAC | |
| GCA GTC TGG ATA AA | TTGG | GGC ACA TCA TGG | TGG GAG CTG G | | | ACTCCCT | ATCGGAGGGATACG | CCCG | CT ACC GAC | |
| GCA GTC TGG | GAG | TCA GGC ACA TCA TGG | TGG GAG CTG G | TTAT | ATGGCATAT | ATTT | TCGGAGGGATACGAG | CTCT | CC GAC | |
| GCA GTC TGG ATA | | GGC ACA TCA TGG | TGG GAG CTG G | CTCC | ATAT | ATCCCAT | ATCGGAGGGATA | AC | GAC | |
| GCA GTC TGG ATA A | CCCGCC | CA GGC ACA TCA TGG | TGG GAG CT | TTCC | | T | ATCGGAGGGATACGAG | GG | CT ACC GAC | |
| GCA GTC TGG ATA A | TTGGA | TCA GGC ACA TCA TGG | TGG GAG CT | TAT | ATGGC | CCCCTCCC | ATCGGAGGGATACGAG | TC | GAC | |
| GCA GTC TGG AT | TC | CA GGC ACA TCA TGG | TGG GAG C | CCCGAT | ATGGC | CCC | ATCGGAGGGATACGAG | | CC GAC | |
| GCA GTC TGG AT | GGG | A GGC ACA TCA TGG | TGG GAG CT | CTAT | | | ATCGGAGGGATACGAG | | CC GAC | |
| GCA GTC TGG | TCCCGA | TCA GGC ACA TCA TGG | TGG GAG CTG G | AGA | | | TCGGAGGGATACGAG | | CT ACC GAC | |
| GCA GTC TGG | TCTGGG | GGC ACA TCA TGG | TGG GAG CT | | | | TCGGAGGGAT | GCT | GAC | |
| GCA GTC TGG A | GA | TCA GGC ACA TCA TGG | TGG GAG CTG G | C | ATGGC | AT | ATCGGAGGGATACGAG | | CT ACC GAC | |
| GCA GTC TGG ATA A | CCCGCC | CA GGC ACA TCA TGG | TGG GAG CTG | | ATGGCATAT | T | ATCGGAGGGAT | G | CC GAC | |
| GCA GTC TGG | | TCA GGC ACA TCA TGG | TGG GAG C | GGGGGGC | ATAT | CCCCCCC | ATCGGAGGGATACGAG | AGGA | ACC GAC | |
| GCA GTC TGG ATA | GGGCGA | TCA GGC ACA TCA TGG | TGG GAG CTG | | | | CGGAGGGATA | AAA | CT ACC GAC | |
| GCA GTC T | CG | TCA GGC ACA TCA TGG | TGG GAG CTG | | | | | A | CT ACC GAC | |
| GCA GTC TGG AT | GGGATCCTCA | TCA GGC ACA TCA TGG | TGG GAG CTG G | | ATGG | A | GGAGGGATACGAG | CCT | CC GAC | |
| GCA GTC TGG | GTA | TCA GGC ACA TCA TGG | TGG GAG CTG | | ATGGC | TCGCAT | ATCGGAGGGATACGAG | CTTG | CC GAC | |
| GCA GTC TGG AT | GGGA | GGC ACA TCA TGG | TGG GAG CT | TA | AT | | ATCGGAGGGATAC | CT | ACC GAC | |
| GCA GTC | GGA | TCA GGC ACA TCA TGG | TGG GAG C | CAG | AT | | ATCGGAGGGATACGAG | GCG | CT ACC GAC | |
| GCA GTC TGG ATA AA | T | GGC ACA TCA TGG | TGG GAG C | AGAT | ATGGCAT | | ATCGGAGGGAT | CGAA | ACC GAC | |
| GCA GTC | CGA | TCA GGC ACA TCA TGG | TGG GAG CTG G | CA | | T | ATCGGAGGGATACGAG | CTCC | ACC GAC | |
| GCA GTC TGG | TCA | TCA GGC ACA TCA TGG | TGG GAG | | | | CGGAGGGATAC | | T ACC GAC | |
| GCA GTC TGG | CCCCTC | TCA GGC ACA TCA TGG | TGG GAG CTG G | CG | | | TCGGAGGGATA | TGGC | GAC | |
| GCA GTC TGG | CCT | TCA GGC ACA TCA TGG | TGG GAG CTG G | CC | AT | | ATCGGAGGGATAC | | GAC | |
| GCA GTC | CGG | TCA GGC ACA TCA TGG | TGG | CCCC | TA | | ATCGGAGGGA | AA | ACC GAC | |
| GCA GTC TGG AT | T | GGC ACA TCA TGG | TGG GAG CT | CCC | GG | | ATCGGAGGGATACGAG | CTGGG | CT ACC GAC | |
| GCA GTC T | TA | TCA GGC ACA TCA TGG | TGG GAG CTG G | A | GG | | GGAGGGATACGAG | GG | CC GAC | |
| GCA GTC TGG | GGA | TCA GGC ACA TCA TGG | TGG GAG CTG G | A | GG | G | CGGAGGGATACGAG | | CT ACC GAC | |
| GCA GTC TGG AT | CGGC | GGC ACA TCA TGG | TGG GAG C | GCAT | ATGGCA | AAAT | ATCGGAGGGATACGAG | CAC | CC GAC | |
| GCA GTC TGG ATA | C | CA GGC ACA TCA TGG | TGG GAG C | ACC | AT | | ATCGGAGGG | G | CT ACC GAC | |
| GCA GTC TGG AT | G | TCA GGC ACA TCA TGG | TGG GAG C | CC | AT | | ATCGGAGGGATACGAG | CTGGGC | GAC | |
| GCA GTC TGG AT | TG | CA GGC ACA TCA TGG | TGG GAG C | CTT | AT | | ATCGGAGGGATACGAG | | CT ACC GAC | |
| GCA GTC TGG | GGA | TCA GGC ACA TCA TGG | TGG GAG CT | CCAT | ATGGCATAT | | ATCGGAGGGATACGAG | | CT ACC GAC | |
| GCA GTC TGG | GGA | TCA GGC ACA TCA TGG | TGG GAG CTG | CTAT | ATGGCA | CG | GGGATACG | | T ACC GAC | |

40

ID3KO (CD24Hi) Singlecel PCR

| Vγ1.1 | P/N | Jγ4 | Vδ6.3 | P/N | Dδ1 | P/N | Dδ2 | P/N | Jδ1 | |
|---|---|---|---|---|---|---|---|---|---|---|
| GCA GTC TGG ATA AA | | TCA GGC ACA TCA TGG | TGG GAG CTG G | | ATGGCATAT | | ATCGGAGGGATACGAG | | CT ACC GAC | Gemline |
| GCA GTC TGG ATA | | GGC ACA TCA TGG | TGG GAG C | | | AT | ATCGGAGGGATACGAG | | CT ACC GAC | (6) |
| GCA GTC TGG | | TCA GGC ACA TCA TGG | TGG GAG C | | | AT | ATCGGAGGGATACGAG | | CT ACC GAC | (3) |
| GCA GTC TGG AT | C | TCA GGC ACA TCA TGG | TGG GAG CTG G | | ATGGCATAT | | TCGGAGGGATACGAG | | CT ACC GAC | |
| GCA GTC TGG AT | C | TCA GGC ACA TCA TGG | TGG GAG CTG | | ATGGCATAT | | TCGGAGGGAT | GC | ACC GAC | |
| GCA GTC TGG | GCCGGA | TCA GGC ACA TCA TGG | TGG GAG CTG G | AGACA | ATAT | AC | AGGG | TCG | CT ACC GAC | |
| GCA GTC TGG | TCAGCA | TCA GGC ACA TCA TGG | TGG GAG CTG | AG | GGCA | CCC | GGGATACGAG | TGG | CC GAC | |
| GCA GTC TGG ATA | TTG | GGC ACA TCA TGG | TGG GAG | AT | ATGGCATAT | A | ATCGGAGGGATACGAG | | CT ACC GAC | |
| GCA GTC TGG | CCC | TCA GGC ACA TCA TGG | TGG GAG CTG G | TAGGG | | AGGG GTT | TCGGAGGGATACG | TGGC | GAC | |
| GCA GTC TGG AT | GACG | GGC ACA TCA TGG | TGG GAG CTG G | | | | GGGATAC | TG | CT ACC GAC | |
| GCA GTC TGG ATA | GGC | GGC ACA TCA TGG | TGG GAG C | CGAT | ATGGCATAT | A | GGAGGGATA | | ACC GAC | |
| GCA GTC TGG ATA | GGC | GGC ACA TCA TGG | TGG GAG CTG G | C | AT | CCCCTCTAT | ATCGGAGGGATACGAG | G | ACC GAC | |
| GCA GTC TGG | GGA | TCA GGC ACA TCA TGG | TGG GAG CT | | | | ATCGGAGGG | TG | CT ACC GAC | |
| GCA GTC TGG ATA | GGC | GGC ACA TCA TGG | TGG GAG CTG G | | TGG | GA | CGGAGGG | TCA | CT ACC GAC | |
| GCA GTC TGG AT | GACG | GGC ACA TCA TGG | TGG GAG C | | | AT | ATCGGAGGGATACGAG | | CT ACC GAC | |
| GCA GTC TGG AT | GGGG | GGC ACA TCA TGG | TGG GAG CTG G | | | T | TCGGAGGGATAC | CTCCTCTGGAC | CT ACC GAC | |
| GCA GTC TGG | TGC | GGC ACA TCA TGG | TGG GAG CT | | | T | ATCGGAGGGATACGAG | | CT ACC GAC | |
| GCA GTC TGG ATA | ACC | TCA GGC ACA TCA TGG | TGG GAG CT | | | T | ATCGGAGGGATACGAG | | CT ACC GAC | |
| GCA GTC TGG A | CC | TCA GGC ACA TCA TGG | TGG GAG CT | | | T | ATCGGAGGGATACGAG | | CT ACC GAC | |
| GCA GTC TGG | GGA | GGC ACA TCA TGG | TGG GAG CTG | | | | ATCGGAGGGATACGAG | CTC | CC GAC | |
| GCA GTC TGG | ACA | GGC ACA TCA TGG | TGG GAG C | | | AT | ATCGGAGGGATACGAG | | CT ACC GAC | |
| GCA GTC TGG | CCG | TCA GGC ACA TCA TGG | TGG GAG CT | | | T | ATCGGAGGGATACGAG | | CT ACC GAC | |
| GCA GTC TGG | CCG | TCA GGC ACA TCA TGG | TGG GAG CTG G | | | AGA | CGGAGGGATACGAG | CTGGGG | GAC | |
| GCA GTC TGG AT | C | GGC ACA TCA TGG | TGG GAG CTG | | | | ATCGGAGGGATACGAG | | CT ACC GAC | |
| GCA GTC TGG ATA | CG | A GGC ACA TCA TGG | TGG GAG CTG G | C | | | ATCGGAGGGATACG | TG | CT ACC GAC | |

30

ID3 WT (CD24LOW) Singlecel PCR

| Vγ1.1 | P/N | Jγ4 | Vδ6.3 | P/N | Dδ1 | P/N | Dδ2 | P/N | Jδ1 | |
|---|---|---|---|---|---|---|---|---|---|---|
| GCA GTC TGG ATA AA | | TCA GGC ACA TCA TGG | TGG GAG CTG G | | ATGGCATAT | | ATCGGAGGGATACGAG | | CT ACC GAC | Gemline |
| GCA GTC TGG ATA | | GGC ACA TCA TGG | TGG GAG CT | | TGG | | CGGAGGGATACGAG | | CT ACC GAC | (2) |
| GCA GTC TGG ATA | | GGC ACA TCA TGG | TGG GAG C | | | AT | ATCGGAGGGATACGAG | | CT ACC GAC | (3) "KO" TCR |
| GCA GTC TGG | | GGC ACA TCA TGG | TGG GAG C | | | AT | ATCGGAGGGATACGAG | | CT ACC GAC | |
| GCA GTC TGG AT | GACG | GGC ACA TCA TGG | TGG GAG C | | | | ATCGGAGGGATACGA | | CT ACC GAC | |
| GCA GTC TGG | | GGC ACA TCA TGG | TGG GAG CT | | TGG | | CGGAGGGATACGAG | | CT ACC GAC | |
| GCA GTC TGG | | GGC ACA TCA TGG | TGG GAG CTG | | | GAT | ATCGGAGGG | | ACC GAC | |
| GCA GTC TGG A | CA | GGC ACA TCA TGG | TGG GAG CTG | | ATG | | TCGGAGGGATACG | GGAGA | ACC GAC | |
| GCA GTC TGG | | GGC ACA TCA TGG | TGG GAG CT | | | T | ATCGGAGGGATACGAG | | CT ACC GAC | |
| GCA GTC TGG A | GA | TCA GGC ACA TCA TGG | TGG GAG CT | | | | TCGGAGGGATACGAG | CTG | T ACC GAC | |
| GCA GTC TGG A | AA | TCA GGC ACA TCA TGG | TGG GAG CTG G | | | | TCGGAGGGATACGAG | CTGTC | GAC | |
| GCA GTC TGG | | GGC ACA TCA TGG | TGG GAG CT | | | T | ATCGGAGGGATACGAG | | CT ACC GAC | |
| GCA GTC TGG | GCA | TCA GGC ACA TCA TGG | TGG GAG CTG | | | | ATCGGAGGGATACGAG | | CT ACC GAC | WT TCR |
| GCA GTC TGG AT | G | GGC ACA TCA TGG | TGG GAG C | | | AT | ATCGGAGGGATACGAG | | CT ACC GAC | |
| GCA GTC TGG | | TCA GGC ACA TCA TGG | TGG GAG CTG G | | | C | CGGAGGGATACGAG | | CT ACC GAC | |
| GCA GTC TGG ATA A | GACGAC | CA GGC ACA TCA TGG | TGG GAG CTG G | | ATGGC | | | TTCCCCTT | T ACC GAC | |
| GCA GTC TGG | GCCAGA | TCA GGC ACA TCA TGG | TGG GAG C | CGC | AT | | ATCGGAGGGATACGAG | CTGTCC | CC GAC | |
| GCA GTC TGG A | CA | TCA GGC ACA TCA TGG | TGG GAG CTG G | CGACTC | | | GAGGGAT | CCCCG | CT ACC GAC | |
| GCA GTC TGG ATA AA | GGTAGAGCGA | TCA GGC ACA TCA TGG | TGG GAG C | GTGT | | | CGGAGGGATACGAG | | CT ACC GAC | |
| GCA GTC TGG A | AA | TCA GGC ACA TCA TGG | TGG GAG CTG G | CTA | | | GGGAT | GCAG | CT ACC GAC | |
| GCA GTC TGG A | GA | TCA GGC ACA TCA TGG | TGG GAG CTG | A | AT | | ATCGGAGGGATAC | CCCCGA | CT ACC GAC | |
| GCA GTC TGG ATA | C | CA GGC ACA TCA TGG | TGG GAG CT | | | A | ATCGGAGGGATACGAG | GA | ACC GAC | |
| GCA GTC TGG | | TCA GGC ACA TCA TGG | TGG GAG CT | | | T | ATCGGAGGGATACGAG | CTC | CC GAC | |
| GCA GTC TGG | GGA | TCA GGC ACA TCA TGG | TGG GAG CTG | | | AC | CGGAGGGATACGAG | | CT ACC GAC | |
| GCA GTC TGG | GGA | TCA GGC ACA TCA TGG | TGG GAG CT | CC | AT | | ATCGGAGGGATACGAG | CCC | CC GAC | |
| GCA GTC TGG AT | TC | CA GGC ACA TCA TGG | TGG GAG CTG G | | | TG | GGAGGGATACGAG | CTG | CC GAC | |
| GCA GTC T | CCCGGGGA | TCA GGC ACA TCA TGG | TGG GAG CTG | AAT | ATGGCATAT | | ATCGGAGGGATACGAG | | CT ACC GAC | |
| GCA GTC TGG | TCCAAA | TCA GGC ACA TCA TGG | TGG GAG CTG G | AAGGGG | | | TCGGAGGGATACG | GGGG G | CT ACC GAC | |
| GCA GTC TGG | GTA | TCA GGC ACA TCA TGG | TGG GAG CTG G | | | GAAG | | TCCCCCCT | CT ACC GAC | |
| GCA GTC TGG ATA | G | CA GGC ACA TCA TGG | TGG GAG CT | AAT | ATGGC | GCAAT | ATCGGAGGGATACGAG | CTTGG | GAC | |
| GCA GTC TGG A | GA | TCA GGC ACA TCA TGG | TGG GAG CTG G | CCT | ATGGCAT | | ATCGGAGGGAT | CCG | CT ACC GAC | |
| GCA GTC TGG ATA | GG | A GGC ACA TCA TGG | TGG GAG CTG G | | | C | CGGAGGGATAC | TCT | CT ACC GAC | |
| GCA GTC TGG | GGA | TCA GGC ACA TCA TGG | TGG GAG | GCC | AT | | ATCGGA | TG | CT ACC GAC | |
| GCA GTC TGG | | TCA GGC ACA TCA TGG | TGG GAG C | | | AT | ATCGGAGGGATACGAG | | CT ACC GAC | |
| GCA GTC TGG A | GA | GGC ACA TCA TGG | TGG GAG CTG G | | | CA | GGAGGGATACGAG | | CT ACC GAC | |

37

ID3KO (CD24LOW) Singlecel PCR

| Vγ1.1 | P/N | Jγ4 | Vδ6.3 | P/N | Dδ1 | P/N | Dδ2 | P/N | Jδ1 | |
|---|---|---|---|---|---|---|---|---|---|---|
| GCA GTC TGG ATA AA | | TCA GGC ACA TCA TGG | TGG GAG CTG G | | ATGGCATAT | | ATCGGAGGGATACGAG | | CT ACC GAC | Gemline |
| GCA GTC TGG | ACC | TCA GGC ACA TCA TGG | TGG GAG CT | | | T | ATCGGAGGGATACGAG | | CT ACC GAC | (10) |
| GCA GTC TGG | | GGC ACA TCA TGG | TGG GAG C | | | AT | ATCGGAGGGATACGAG | | CT ACC GAC | (6) |
| GCA GTC TGG ATA | | GGC ACA TCA TGG | TGG GAG CT | | | T | ATCGGAGGGATACGAG | | CT ACC GAC | (5) |
| GCA GTC TGG | | GGC ACA TCA TGG | TGG GAG CTG G | | | | TCGGAGGGATACGAG | | CT ACC GAC | (3) |
| GCA GTC TGG ATA | | GGC ACA TCA TGG | TGG GAG C | | | AT | ATCGGAGGGATACGAG | | CT ACC GAC | (3) "KO" TCR |
| GCA GTC TGG | | GGC ACA TCA TGG | TGG GAG | | | T | GGAGGGATACGA | | CT ACC GAC | |
| GCA GTC TGG | | TCA GGC ACA TCA TGG | TGG GAG | | | T | GGAGGGATACGA | | CT ACC GAC | |
| GCA GTC TGG | | TCA GGC ACA TCA TGG | TGG GAG CTG | | | | ATCGGAGGGATACGA | C | CT ACC GAC | |

30

**Figure S2. Effect of Id3 deficiency on the diversity of CDR3γ and CDR3δ diversity of the Vγ1.1Vδ6.3 TCRs.**

Single cell sequencing was performed for the V regions of the Vγ1.1 and Vδ6.3 subunits of $CD24^{hi}$ immature and $CD24^{low}$ mature γδ T cells from *Id3*$^{+/+}$ and *Id3*$^{-/-}$ mice. The sequences of the clonotypes are listed relative to germline sequence, and their representation relative to the total number of clonotypes sequences is indicated in parenthesis on the right. The clonotype sequence of the Vγ1.1Vδ6.3 TCRs used to make the WT and KO Tg mice are highlighted in yellow.

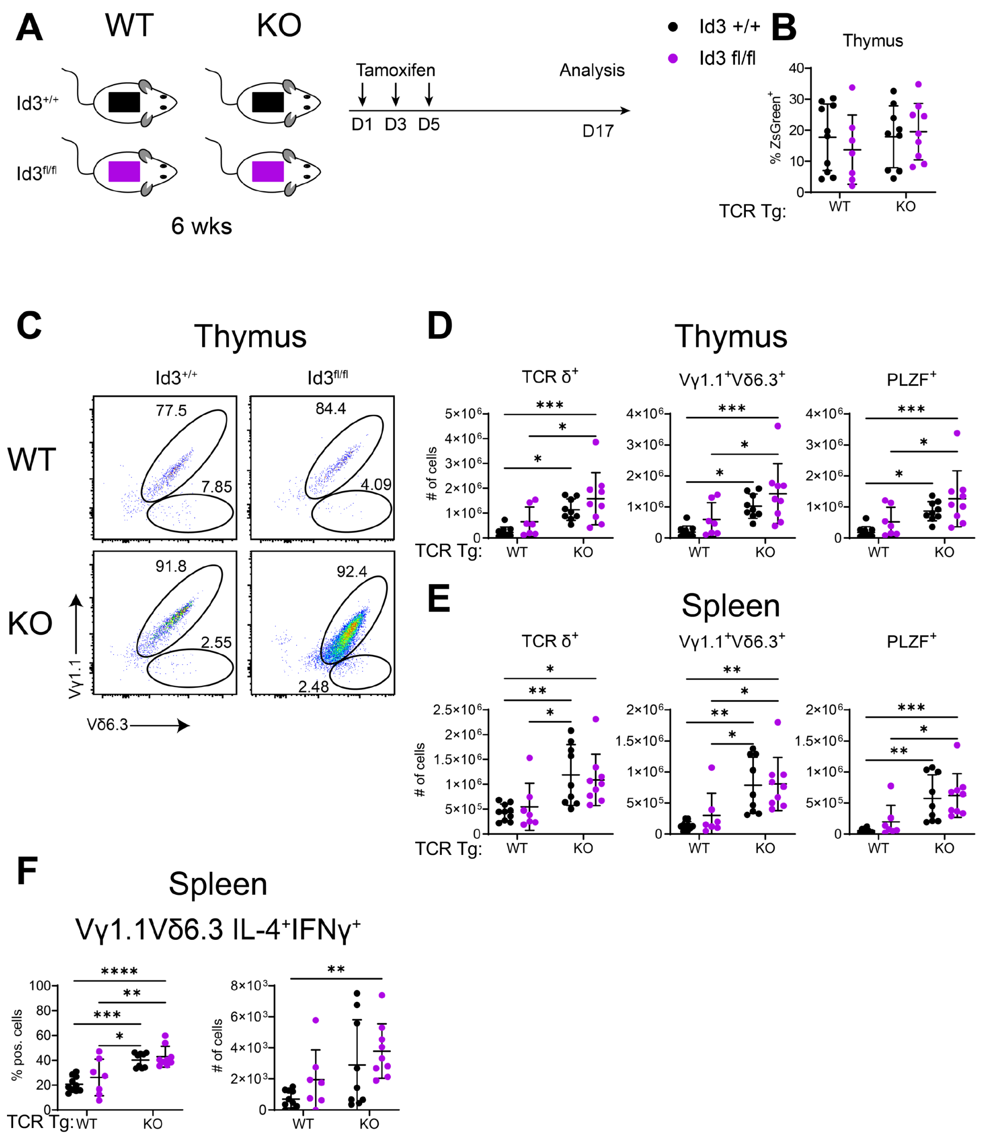


**Figure S3. Impact of conditional ablation of *Id3* using TCRd-Cre on development of Vγ1.1Vδ6.3 TCR expressing NKγδT cells.**

(A) Diagram of *in vivo* experimental design. Tamoxifen was administered to 6-week-old mice on days 1, 3, and 5, and analysis was performed on day 17.

(B) Graphic depiction of the frequency of ZsGreen$^+$ thymocytes.

(C) Representative flow cytometry plots of ZsGreen$^+$TCRδ$^+$ thymocytes displayed as Vγ1.1 vs Vδ6.3 for mice expressing the WT or KO Vγ1.1Vδ6.3 TCR.

(D,E) Number of thymic (D) or splenic (E) TCRδ$^+$, Vγ1.1$^+$Vδ6.3$^+$, and PLZF$^+$ Vγ1.1$^+$Vδ6.3$^+$ cells is depicted graphically with each symbol representing a single mouse.

(F) Graphic representation of the frequency and number of IL-4 and IFNγ co-producing Vγ1.1$^+$Vδ6.3$^+$ splenocytes following PMA/Ionomycin stimulation, with each symbol representing a single mouse.

$TCR^{WT}Id3^{+/+}$ (n = 10), $TCR^{WT}Id3^{fl/fl}$ (n = 7), $TCR^{KO}Id3^{+/+}$ (n = 9), and $TCR^{KO}Id3^{fl/fl}$ (n = 9). Data were pooled from 3 independent experiments and are plotted as mean±SD. Statistical analysis: two-way ANOVA with correction for multiple comparison using Tukey's post hoc testing *$p < 0.05$, **$p < 0.01$, ***$p < 0.001$, ****$p < 0.0001$.

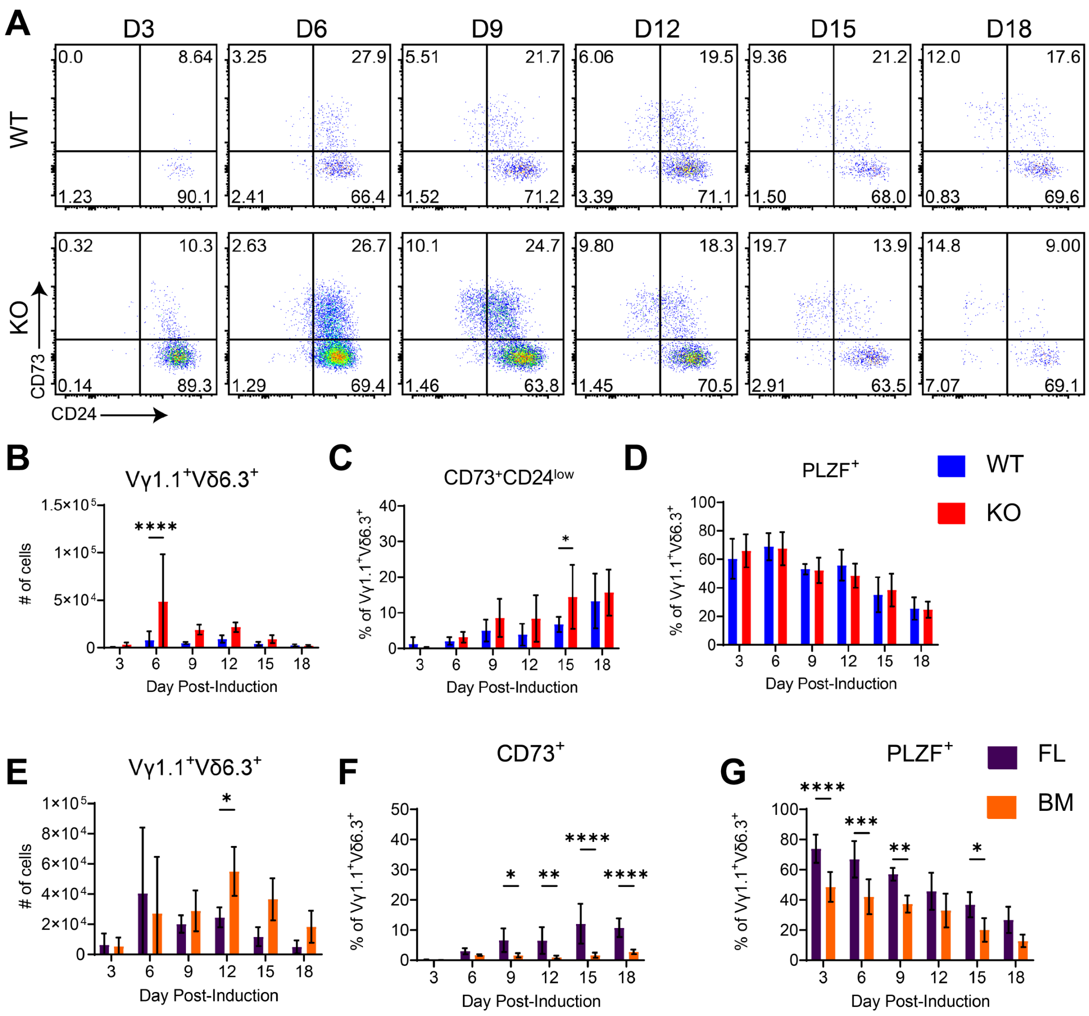


**Figure S4. Capacity of Vγ1.1Vδ6.3 TCR complexes to instruct the NKγδT cell fate in vitro.**

(A) Representative flow cytometry plots of FL derived Vγ1.1Vδ6.3 T lymphocyte maturation in culture displayed as CD73 vs CD24. The WT and KO Tg TCRs were induced in FL derived precursors cultured in T cell polarizing conditions.

(B-D) Graphic depiction of the (B) absolute numbers Vγ1.1Vδ6.3 expressing cells obtained, (C) frequency which were $CD73^{+}CD24^{low}$, and (D) frequency of Vγ1.1Vδ6.3 cells expressing PLZF.

(E-G) The KO Tg TCR was induced in FL and bone marrow derived precursors cultured in T cell polarizing conditions. Graphic depiction of the (E) absolute numbers Vγ1.1Vδ6.3 expressing cells obtained, and the frequencies of these which were (F) $CD73^{+}CD24^{low}$ or (G) expressed PLZF.

Data are pooled from 3 independent experiments (n = 6). Data are plotted as mean±SD. Statistical analysis: two-way ANOVA with correction for multiple comparison using Sidak's multiple comparisons test. *$p < 0.05$, **$p < 0.01$, ***$p < 0.001$, ****$p < 0.0001$.

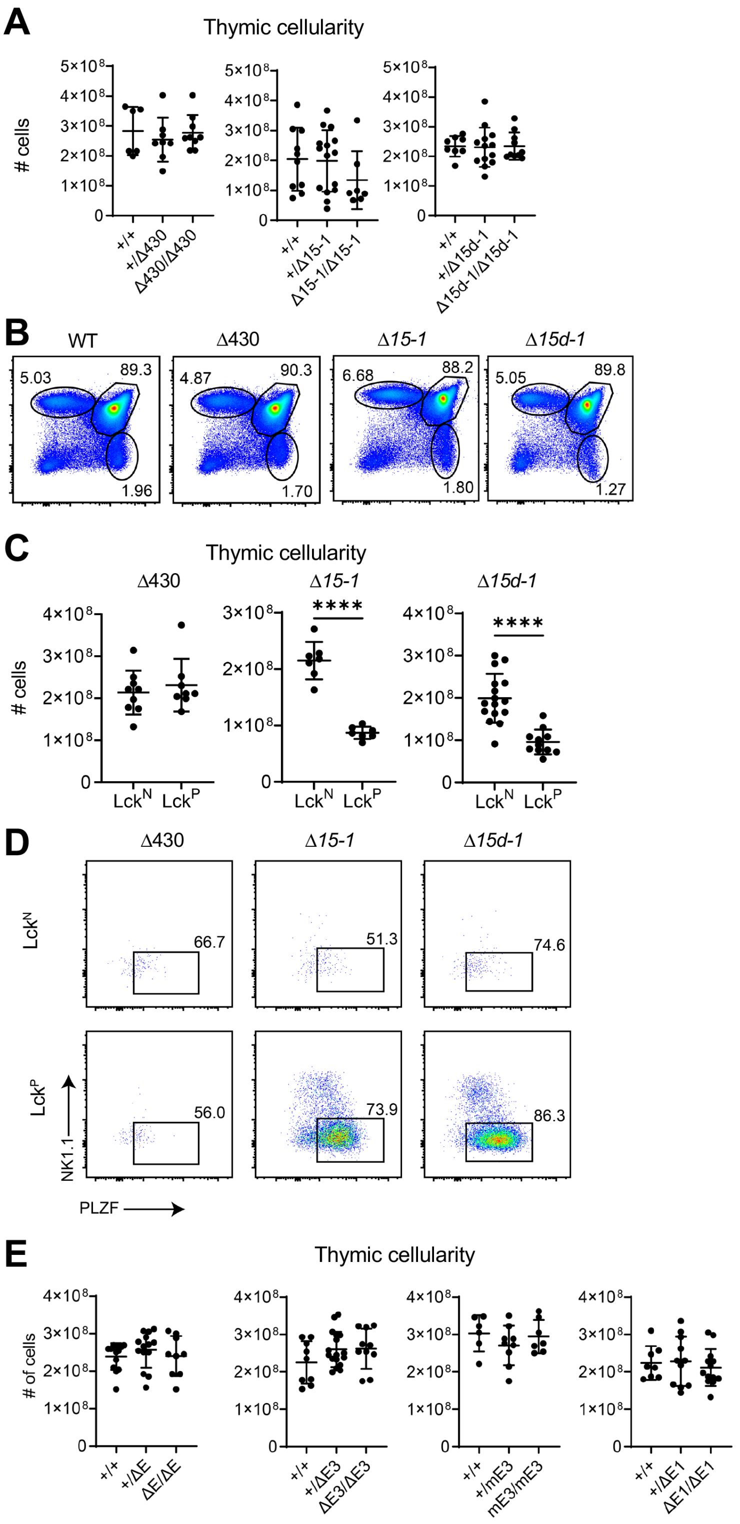


**Figure S5. Role of E protein binding in *Trav15* element selection in developing NKγδT cells.** (A) Thymic cellularity in *Δ430, Δ15-1*, and *Δ15d-1* mutant mice is depicted graphically with each symbol representing an individual mouse. All comparisons involved littermates. *Δ15-1: Tcra*$^{+/+}$ (n = 10), *Tcra*$^{+/\Delta 15\text{-}1}$ (n = 15), and *Tcra*$^{\Delta 15\text{-}1/\Delta 15\text{-}1}$ (n = 7); *Δ15d-1: Tcra*$^{+/+}$ (n = 8), *Tcra*$^{+/\Delta 15d\text{-}1}$ (n =

13), and *Tcra*$^{\Delta 15d\text{-}1/\Delta 15d\text{-}1}$ (n = 10); *Δ430: Tcra*$^{+/+}$ (n = 6), *Tcra*$^{+/\Delta 430}$ (n = 8), and *Tcra*$^{\Delta 430/\Delta 430}$ (n = 9). Data were pooled from a minimum of 3 independent experiments and are plotted as mean±SD. Statistical analysis: one-way ANOVA with correction for multiple comparison using Tukey's post hoc testing. Significant differences were not detected.

(B) Representative flow cytometry plots for subsets defined by CD4 and CD8 expression in *Tcra*$^{+/+}$*, Tcra*$^{\Delta 430/\Delta 430}$*, Tcra*$^{\Delta 15\text{-}1/\Delta 15\text{-}1}$*, Tcra*$^{\Delta 15d\text{-}1/\Delta 15d\text{-}1}$ mutant mice.

(C) Thymic cellularity in Lck-Cre negative (Lck$^{N}$) Id3 sufficient (*Id3*$^{fl/fl}$) (control), or Lck-Cre (Lck$^{P}$) mediated Id3 deficient (*Id3*$^{fl/fl}$) *Δ430, Δ15-1,* and *Δ15d-1* mutants. All comparisons to Lck-Cre negative (Lck$^{N}$) littermates. For *Δ430:* Lck$^{N}$ (n = 9), Lck$^{P}$ (n = 8); *Δ15-1*: Lck$^{N}$ (n = 7), Lck$^{P}$ (n = 7); *Δ15d-1:* Lck$^{N}$ (n = 16) and Lck$^{P}$ (n = 11). Data are pooled from at least 3 independent experiments and plotted as mean±SD. Statistical analysis: Student's t test. ****$p < 0.0001$.

(D) Representative flow cytometry plots of NK1.1 and PLZF expression in Vγ1$^{+}$ thymocytes from Lck-Cre negative (Lck$^{N}$) Id3 sufficient (*Id3*$^{fl/fl}$) (control), or Lck-Cre (Lck$^{P}$) mediated Id3 deficient (*Id3*$^{fl/fl}$) *Δ430, Δ15-1,* and *Δ15d-1* mutants

(E) Thymic cellularity in *ΔE, ΔE3, mE3,* and *ΔE1* mutant mice is depicted graphically with each symbol representing an individual mouse. All comparisons involved littermates. *ΔE: Tcra*$^{+/+}$ (n = 14), *Tcra*$^{+/\Delta E}$ (n = 12), and *Tcra*$^{\Delta E/\Delta E}$ (n = 9); *ΔE3: Tcra*$^{+/+}$ (n = 8), *Tcra*$^{+/\Delta E3}$ (n = 17), and *Tcra*$^{\Delta E3/\Delta E3}$ (n = 10); *mE3: Tcra*$^{+/+}$ (n = 6), *Tcra*$^{+/mE3}$ (n = 9), and *Tcra*$^{mE3/mE3}$ (n = 7); *ΔE1: Tcra*$^{+/+}$ (n = 8), *Tcra*$^{+/\Delta E1}$ (n = 11), and *Tcra*$^{\Delta E1/\Delta E1}$ (n = 13). Data were pooled from a minimum of 3 independent experiments and are plotted as mean±SD. Statistical analysis: one-way ANOVA with correction for multiple comparison using Tukey's post hoc testing. Significant differences were not detected.
